# Retrosplenial cortex enables context-dependent goal-directed sensorimotor transformation

**DOI:** 10.1101/2025.10.31.685738

**Authors:** Pol Bech, Robin F. Dard, Jules Lebert, Lana Smith, Axel Bisi, Anthony Renard, Sylvain Crochet, Carl C.H. Petersen

## Abstract

The ability to dynamically adjust a behavioral response to a stimulus depending on context is of critical importance for animals. To investigate the neural basis supporting context-dependent sensory processing we developed a behavioral task in which mice changed their response to a single whisker deflection according to a continuously present contextual cue. Through unbiased optogenetic inactivation mapping, we found that neuronal activity in sensory and motor cortices contributed to task execution and, interestingly, we uncovered an unexpected role of retrosplenial cortex for contextual integration. Widefield calcium imaging revealed that retrosplenial cortex was the first dorsal cortical area to show context discrimination in response to whisker stimulation, followed by whisker motor cortex. Finally, we combined optogenetic inactivation with calcium imaging to define causal context-dependent changes in sensorimotor processing. Our cortex-wide mapping experiments thus begin to define key cortical nodes for context-dependent sensorimotor transformation and highlight an important contribution of retrosplenial cortex.

## Introduction

In order to choose the appropriate action in response to a given sensory stimulus, animals typically need to integrate surrounding “contextual” information. While we begin to understand mechanisms underlying early sensory processing and the transformation of sensations into learned actions (Chen et al., 2024, 2015, 2017; Curtis and Kleinfeld, 2009; Esmaeili et al., 2021; Guo et al., 2014; Harris and Mrsic-Flogel, 2013; International Brain Laboratory et al., 2025; Khilkevich et al., 2024; Kim et al., 2021; Kleinfeld and Deschênes, 2011; Kwon et al., 2016; Mayrhofer et al., 2019; O’Connor et al., 2010; Resulaj et al., 2018; Sachidhanandam et al., 2013; Siegle et al., 2021; Yang et al., 2016), less is known regarding how functionally coupled sensorimotor associations can be dynamically modified by context (Allen et al., 2019; Banerjee et al., 2020; Chang et al., 2024; Chevée et al., 2022; Cole et al., 2024; Condylis et al., 2020; Gershman, 2017; Gilbert and Li, 2013; Kuchibhotla et al., 2017; Ortiz et al., 2020; Rossi-Pool et al., 2021; Wu et al., 2020).

The definition of context is large enough for multiple general categories to emerge. At its most fundamental level, people have often considered as context internal processes or states like motivation level, attention level or ongoing action. Decades of studies in this context modality have identified thalamocortical and neuromodulator-dependent disinhibitory mechanisms responsible for modifying sensory processing and cortical-network states (Crochet and Petersen, 2006; Eggermann et al., 2014; Gasselin et al., 2021; Lohani et al., 2022; Poort et al., 2022; Poulet et al., 2012; Poulet and Petersen, 2008; Saalmann et al., 2012; Urbain et al., 2015; Zhou et al., 2016). A different variation on context, this time considering the influence of external environmental stimuli on perception, learning, and memory, has been extensively studied in contextual fear/appetitive conditioning, navigation/foraging tasks, or working memory tasks. A large body of evidence in contextual fear conditioning shows it relies on well described circuits involving among others, the amygdala, the prelimbic system, the prefrontal cortex and the hippocampus (Tovote et al., 2015). Context-dependent navigation tasks depend on the ability of the hippocampus and associated regions such as the entorhinal cortex or the retrosplenial cortex to represent both spatial and context information. Interactions between the hippocampus and the prefrontal cortex supports context-dependent decision making in an exploration task (Place et al., 2016). Other studies have investigated the mechanisms underlying behavioral flexibility in rule changing paradigms in which context must be inferred by the animal. This ability to infer context and adjust a behavioral response accordingly depends on PFC (Birrell and Brown, 2000; Marton et al., 2018), orbitofrontal cortex (Banerjee et al., 2020; Klein-Flügge et al., 2022; Schoenbaum et al., 2009), premotor regions such as wM2 and ALM (Chang et al., 2024; Finkel et al., 2024), and anterior cingulate cortex (Cole et al., 2024; Hajnal et al., 2024). It is also possible to deconstruct context into a discrete sensory cue, like a pure tone or visual flash, to provide contextual information which modifies the contingency of a second upcoming stimulus. The ability to execute this type of tasks requires the maintenance of the information given by the first, context-setting cue to guide the response to the second cue, which is mediated by working memory. Many studies have shown that working memory is supported by prefrontal and medial prefrontal cortex (Asaad et al., 1998; Duan et al., 2015; Goldman-Rakic, 1995; Hasselmo, 2006; Liu et al., 2014; Romo et al., 1999; Stern et al., 2001). In mice, planification of movement execution relies on the anterolateral motor cortex (ALM), as well as wM2 (Esmaeili et al., 2021; Guo et al., 2014). Recent work has presented evidence that ALM also contributes to context-dependent decisions in a delayed match to sample task (Wu et al., 2020), presumably by disrupting the working memory representations necessary for maintenance of the identity of the sample.

Multiple parallel streams of research have therefore investigated various important brain regions mediating context-dependent behaviors, with experiments conducted in mice increasingly aiming to probe causal circuit mechanisms through applying new advanced methodologies. Whether some of these mechanisms are shared across modalities of context-dependent behaviors remains to be investigated. Mice have a highly-developed array of facial whiskers, which they use to determine the size, shape, location and texture of objects in their immediate environment (Diamond et al., 2008; Feldmeyer et al., 2013; Petersen, 2019; Staiger and Petersen, 2021). Detection of a whisker deflection reported by licking for reward is among the simpler tasks explored to date, but nonetheless evidence suggests that many brain regions contribute to task execution, including various cortical regions, thalamus, striatum, and superior colliculus (El-Boustani et al., 2020; Kwon et al., 2016; Le Merre et al., 2018; Mayrhofer et al., 2019; Miyashita and Feldman, 2013; Oryshchuk et al., 2024; Sachidhanandam et al., 2013; Sippy et al., 2015; Takahashi et al., 2020; Yamashita and Petersen, 2016; Yang et al., 2016). Building on this relatively simple whisker detection task, we now begin to ask if we can adapt it to study specific circuits for contextual modulation of sensory processing. To address this question, here, we developed a context-dependent detection task where the reward contingency of a well-defined single-whisker sensory stimulus was modulated according to the texture of auditory background noise. Combining unbiased cortex-wide inactivation screening with widefield calcium imaging, we identified cortical regions involved in task execution and found that retrosplenial cortex appears to play a key role for integration of sensory stimuli and context.

## Results

### Rapid behavioral adaptation to sensory stimuli in a changing context

We trained head-fixed, water-restricted mice to adapt their response to a whisker sensory stimulus according to an explicitly and continuously given contextual cue in the form of two distinct auditory background sounds (pink noise or brown noise). We deterministically alternated between contexts in blocks of 20 pseudo-randomly organized trials during single sessions. According to this changing context, mice were rewarded (or not) for licking a water spout within a 1-s response window following a brief (3 ms) stimulus of the C2 whisker, whereas licking after a brief (10 ms) auditory 10 kHz tone was always rewarded (Figure 1A). Thus, mice had to learn to lick for reward in response to the whisker stimulus only in the whisker-rewarded (W+) context and refrain from licking in the whisker-non-rewarded (W-) context, while maintaining constant licking in response to the auditory pure tone in both contexts. In addition, no-stimulus trials (Catch) were randomly delivered to assess the spontaneous lick rate. Error licks in Catch trials or whisker stimulus trials in W- blocks were neither rewarded nor punished.

**Figure 1.**
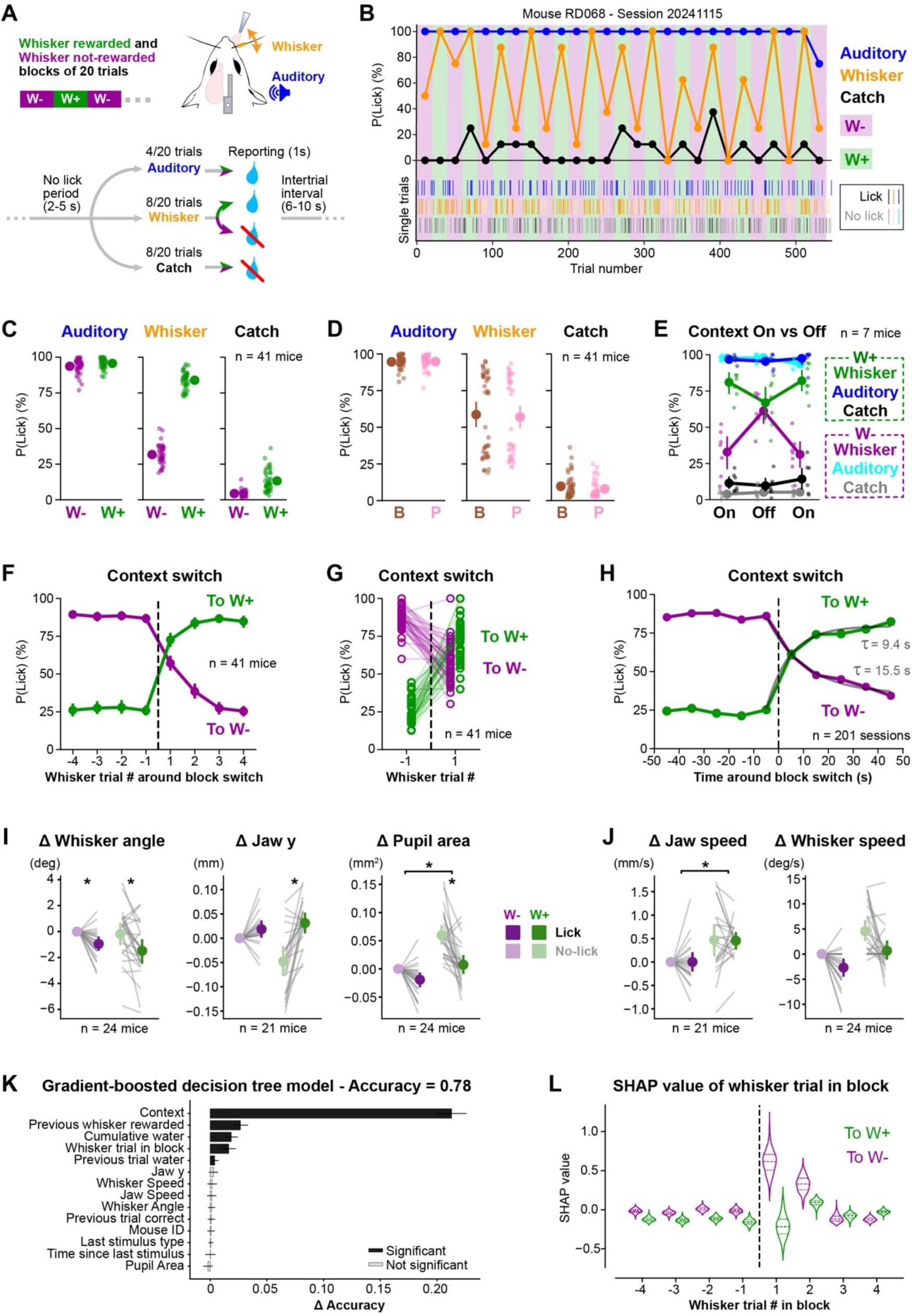
A context-dependent whisker detection task. (**A**) Schematic of the task structure. (**B**) Block-average lick probability in one example expert session for whisker trials (orange), auditory trials (blue) and catch trials (black). W+ and W- blocks are shown by green and purple shaded areas, respectively. Single trials are shown by individual ticks (below). (**C**) Averaged lick probability for whisker, auditory and catch trials in W+ and W- contexts. Small dots represent individual mice (n = 41 mice, 201 sessions); large-filled circles represent grand-averages with error bars showing the 95% confidence interval. (**D**) Same as C, but split according to brown (B) or pink (P) noise context identity. (**E**) As a control, the contextual brown and pink noises (Context On) were replaced by constant white noise (Context Off) for one session (n = 7 mice). (**F**) Averaged lick probability for whisker trials centred on context transitions. From W-to W+ contexts in green and from W+ to W- contexts in purple (n = 41 mice). (**G**) Lick probability at last and first whisker trials around context transition (n = 41 mice). (**H**) Lick probability in response to the last and first whisker trial around context transition according to the delay at which it occurs. Transition from W- to W+ contexts in green and from W+ to W- contexts in magenta. Grey solid lines show exponential fits of the averaged lick probability over time. (**I&J**) Quantification of the changes in orofacial posture (panel I) and movements (panel J) during the no-lick window preceding whisker trial onset, relative to No-lick trials in the W- context. Each line represents an individual mouse; large-filled circles represent grand-averages with error bars showing the 95% confidence interval. (**K**) Drop in accuracy for permutation of labels for each of the parameters of the gradient-boosted model predicting single whisker-trial responses. (**L**) SHAP values for the ‘whisker trial in block’ parameter aligned on context transition from W+ to W- in magenta and from W- to W+ in green.

Mice were trained in this task until they reached expert performance, completing an average of 23 blocks per session (Figure 1B and Figure 1 – figure supplement 1A-E). In expert sessions, mice showed consistent licking responses to the auditory stimulus in both contexts (Auditory hit rate: W+ = 96 ± 4 %, W- = 94 ± 5 %; n = 41 mice), and strong context-modulated licking in response to whisker stimuli (W+ = 84 ± 6 %, W- = 32 ± 7 %) (Figure 1C and Figure 1 – figure supplement 1D), irrespective of which background noise was associated to each context (Figure 1D). Lick rates in catch trials were overall low, although slightly higher in W+ blocks (Catch false alarm rate: W+ = 13 ± 7 %, W-: 4 ± 2%; t-value = -9, p-value = 1.6 x 10^-10^). The consistent lick responses to the auditory pure tone suggest that the modulation of the responses to the whisker stimulus is not due to disengagement. To test if mice used the background contextual information to guide their choices, we compared performance of a subset of mice in the original task (Context On) with a variation in which both context sounds were replaced by continuous white noise while preserving the underlying block reward structure (Context Off). After removing the explicit contextual information mice were unable to discriminate between context blocks (Figure 1E and Figure 1 – figure supplement 1F).

We next investigated how quickly mice adapted their responses to the whisker stimulus following context transition. Mice changed their licking behavior as early as the first whisker trial following the context transition with a slight gradual further adaptation over subsequent whisker trials (Figure 1F). The probability of licking in response to whisker deflection was significantly different comparing the first trial after context change with the last trial of the previous block (transition from W- to W+: pre = 26 ± 8 %, post = 73 ± 13 %, t-value = 21, p-value = 2.8 x 10^-23^; transition from W+ to W-: pre = 87 ± 7 %, post = 57 ± 13 %, t-value = -14, p-value = 7.3 x 10^-17^) (Figure 1G). Since trials within each context block were pseudo-randomly ordered, the first whisker trial could occur at variable delays after the context transition. To assess whether this delay influenced the mouse response, we analysed the lick probability to the first whisker trial in either the W+ or W− context as a function of time elapsed since the transition (Figure 1H). We observed a progressive increase in lick probability in the W+ context and a corresponding decrease in the W− context, suggesting temporal integration of contextual information after context switch. To assess whether this temporal integration would differ between contexts we fitted an exponential to the time evolution of the lick probability. This suggested a faster transition to the W+ context than to the W- context (W+ time constant: 9.4 s, W- time constant: 15.5 s) (Figure 1H). However, there was no apparent sign of anticipation of the context switch in the behavior of the mice.

Context could influence the mouse posture or uninstructed movements, which could reflect active sensing or attentional changes (Musall et al., 2019). To investigate context- or performance-specific behavioral changes in expert sessions, we performed high-speed videography of the mouse face. Using pose estimation (Mathis et al., 2018), we extracted dorsal and lateral orofacial kinematic features, including whisker angle, jaw opening, and pupil area (Figures 1I&J and Figure 1 – figure supplement 2). We first sought to investigate whether the pre-stimulus baseline orofacial posture and motion could reflect the current context and / or could predict a licking response to the upcoming whisker stimulus. We analysed the 2-s no-lick window before whisker trial onsets and computed the average position and speed. Our analysis revealed no significant general effect of context for the average whisker angle or jaw opening, but pupil area was larger in the W+ context, perhaps reflecting attentional state (McGinley et al., 2015; Reimer et al., 2014) (Figure 1I and Figure 1 – figure supplement 2B). Furthermore, the average whisker angle was more posterior and the jaw rested in a more open position in the pre-stimulus period of trials in which the mouse subsequently licked in response to the whisker stimulus independently of context. We also considered whether anticipatory movements of the whiskers and jaw could explain behavioral performance, finding no effect of context or lick reflected in the speed of whisking of the mouse, while jaw speed was significantly increased in the W+ context (Figure 1J and Figure 1 – figure supplement 2C).

We next wanted to assess the combined influence of task parameters such as context, trial index and history, as well as orofacial movement metrics onto task execution and specifically onto choice in response to whisker trials in the W+ and W-contexts. We trained a gradient boosted tree model (Griffiths et al., 2024) (Figures 1K&L and Figure 1 – figure supplement 3) on concatenated expert sessions including only whisker trials and using uncorrelated features. We used SHAP values (Lundberg and Lee, 2017) to obtain single-trial contributions from each parameter to model outcome. SHAP values can be interpreted as the bias that a single parameter inflicts on predicting outcome toward lick or no lick. Our model reached a cross-validated accuracy of 0.78 on held-out test trials (Figure 1K). Among all model features, context information was the one leading to the strongest accuracy drop in the permutation test, followed by whether the previous whisker trial was rewarded or not, the cumulative number of rewards (indicative of the level of thirst motivation), and the index of the whisker trial of the block (Figure 1K). Following context transition, mice adapt their response to whisker stimulation already at the first trial of the block, but further improve their choice on the second whisker trial (Figure 1F). To assess if the model could account for this effect, we examined the SHAP value of the parameter “whisker trial in block”, finding a pattern consistent with the mouse behavior capturing both single trial adaptation after context transition and a slower component (Figure 1L). Overall, the experimental data and the model indicate that the explicit auditory contexts presented in W+ and W- blocks appear to dominate task performance with additional contributions associated with ongoing behavioral state and reward history.

### Distributed cortical areas are involved in task execution

To identify cortical regions involved in task performance, we used an unbiased optogenetic inactivation approach (Guo et al., 2014) using a 1 mm-spaced grid of target locations covering the left hemisphere of the mouse brain. We trained VGAT-ChR2 mice (Zhao et al., 2011) in the context task and, after reaching expert performance, we pseudo-randomly inactivated individual cortical sites in half of the trials by directing a blue light beam to specific grid positions (Figures 2A&B and Figure 2 – figure supplement 1A&B). As a control condition, in the other half of the trials, the beam was directed outside the brain. For each trial type in both W+ and W− contexts, we quantified changes in lick probability relative to these control trials (Figures 2C-E). In catch trials, optogenetic inactivation had little effect on licking behavior in either context (Figure 2C). However, in auditory trials, there was a reduction in lick probability when inactivating auditory cortex and frontal premotor regions, which was similar across contexts (Figure 2D). Finally, in whisker trials, we observed that inactivating primary and secondary whisker somatosensory cortices (wS1 and wS2), as well as anterior premotor regions led to a reduction of lick probability across both contexts (Figure 2E and Figure 2 – figure supplement 1C). Interestingly, during whisker trials, inactivation of the tongue–jaw primary somatosensory cortex (tjS1) and the retrosplenial cortex (RSC) increased lick probability with respect to the control condition specifically in W− trials, indicating a context-dependent effect of these two regions (Figure 2E, right panel). In control mice without ChR2 expression, we did not observe any significant changes in lick probability following optogenetic inactivation at any site (Figure 2 – figure supplement 1D&E). Pharmacological inactivation using muscimol injected into wS1, RSC, or the forepaw primary somatosensory cortex gave results consistent with optogenetic inactivation (Figure 2 – figure supplement 2). While lick latency increased during inactivation of frontal areas, presumably correlated with decreased motor function, there was no change in lick latency associated with inactivation of RSC or tjS1 (Figure 2F and Figure 2 – figure supplement 3). Inhibition of tjS1 or RSC therefore does not seem to directly release a motor response, but rather is more likely to act via downstream brain regions.

**Figure 2.**
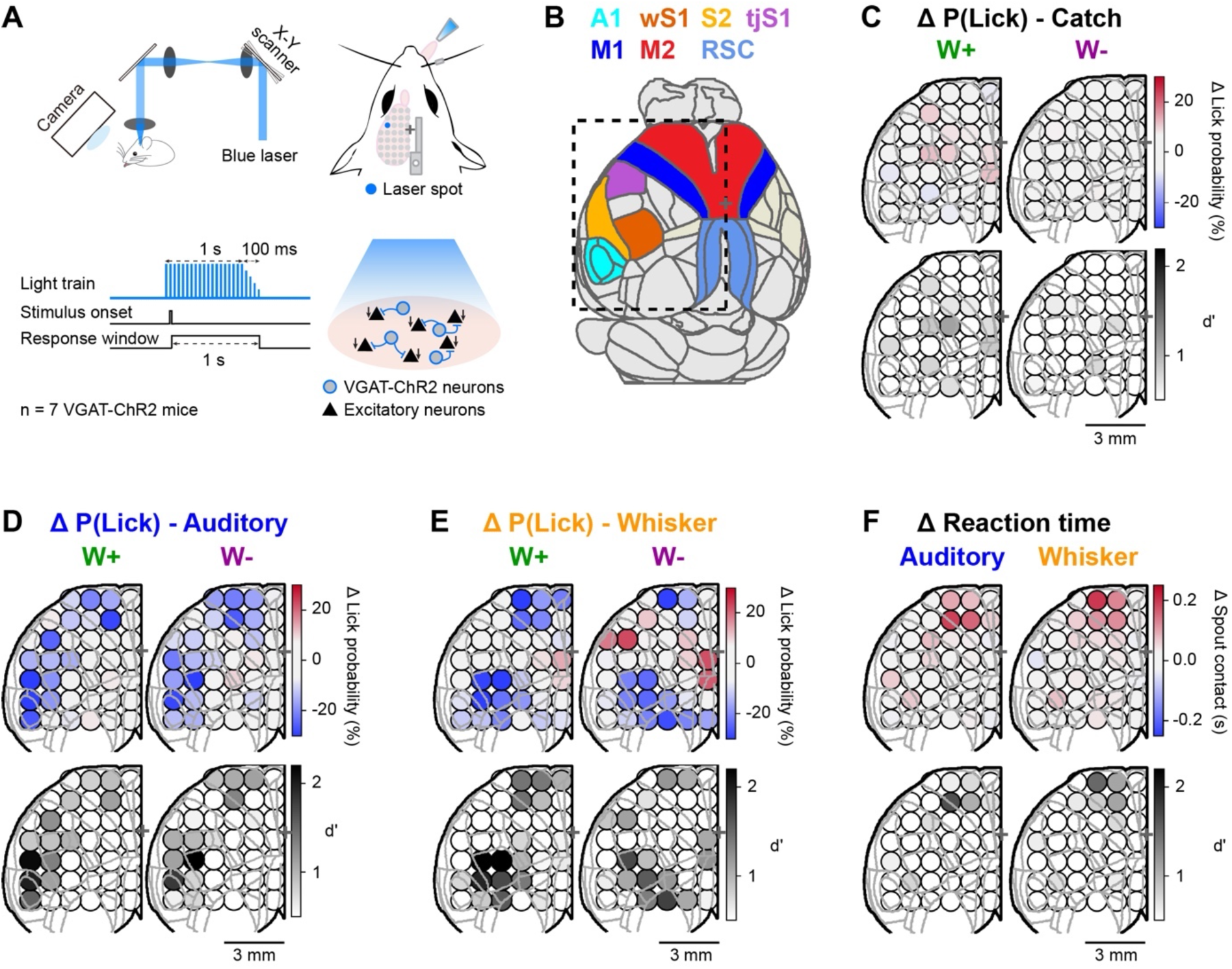
Optogenetic inactivation screen. (**A**) Schematic of the optoinhibition screening in VGAT-ChR2 mice. (**B**) 30° rotated top view of the Allen mouse brain atlas, as in the experimental setup. (**C**) Top: Grid map showing changes in lick probability, ΔP(Lick), evoked by optogenetic inactivation of the target area during catch trials in W-and W+ contexts. Bottom: d’ computed between the distribution of ΔP(Lick) in optoinhibition trials and a null distribution computed by shuffling all trials 1,000 times. (**D**) Same as C, but for auditory trials. (**E**) Same as C, but for whisker trials. (**F**) Top: Grid map showing change in lick reaction time (tongue-spout contact time) in correct response to auditory and whisker trials during optogenetic inactivation of the target area relative to control trials. Bottom: associated d’.

Thus, unbiased optogenetic and targeted pharmacological inactivation revealed several cortical regions critical for the execution of the context task, including wS1, wS2, wM1/2 and ALM, known to be required for whisker-based sensory-to-motor transformations. Furthermore, we identified RSC and tjS1 as key regions which influence the ability of the mouse to adapt its response during whisker trials in a context-dependent manner.

### Context-dependent flow of activity during sensorimotor transformation

To investigate the spatiotemporal neural dynamics underlying task execution, we recorded calcium activity across the dorsal cortex in transgenic mice expressing jRGECO1a (Dana et al., 2018; Tamura et al., 2025) (Figure 3) or GCaMP6f (Daigle et al., 2018; Harris et al., 2014) (Figure 3 – figure supplement 1), obtaining consistent results. We focused our analyses on the first 120 ms after sensory stimuli, largely preceding the onset of jaw opening in whisker trials in the W+ context (103 ± 12 ms) (Figures 3B&C). We then compared the evoked cortical activity in auditory trials across W+ and W- contexts, and observed neural activity patterns dominated by an early increase in fluorescence in the primary auditory cortex (A1), followed by RSC (and nearby regions), and, subsequently, activity propagated anteriorly towards a medial part of secondary motor region (including wM1/2), before transitioning to the orofacial sensorimotor cortices (including tjS1 and ALM) (Figures 3D&E). The auditory evoked responses did not differ comparing W+ and W- contexts. In contrast, whisker trials exhibited context-dependent differences in cortical dynamics (Figures 3F&G). Early after stimulus onset, whisker deflection evoked similar activation of primary and secondary whisker somatosensory cortices (wS1 and wS2) in both W+ and W− contexts. However, after ∼50 ms post-stimulus, cortical dynamics began to diverge, with increased activity in RSC and wM1/2 in the W+ context compared to the W-context. To investigate whether the RSC and wM1/2 modulation was context specific or lick related we compared whisker lick and auditory lick-evoked cortical activity patterns in the W+ context. Besides sensory modality-specific activity such as the early increase of activity in wS1/2 and wM1/2 in whisker trials or A1 in auditory trials (Figure 3 – figure supplement 2), we observed generally stronger activation in RSC in whisker-lick trials in the W+ context compared to auditory-lick trials. Importantly, these dynamics were not observed in control mice expressing tdTomato or GFP (Figure 3 – figure supplement 1).

**Figure 3.**
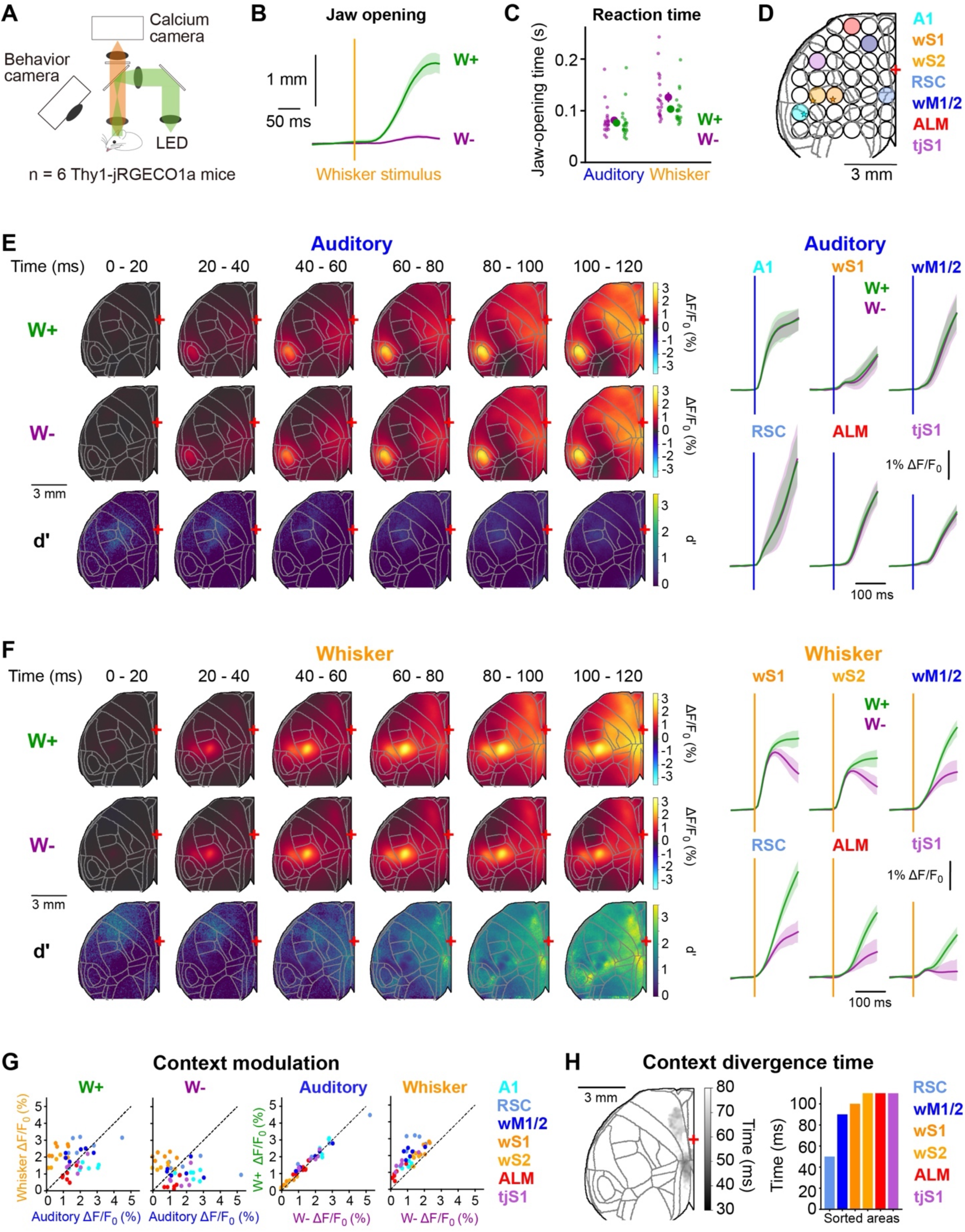
Imaging cortical spatiotemporal dynamics of context-dependent sensorimotor processing. (**A**) Experimental apparatus. (**B**) Averaged time course of jaw opening aligned on whisker stimulus for correct trials in W+ (green) and W- (purple) contexts. (**C**) Reaction time in auditory and whisker trials in W- and W+ contexts derived from the time course of the jaw opening. (**D**) Grid map showing the target areas that are used as ROIs for trial averaged activity. The stars represent the centre of mass of manually drawn ROIs centred on the peak of sensory-evoked activity for each mouse (auditory trials for A1; whisker trials for wS1 and wS2). (**E**) Left: Image series showing evolution of cortical dynamics following auditory trials in the W+ context (first row), in the W- context (second row), and the time-by-time distance (d’) between W-and W+ context activity maps (third row). Right: Average time-course of stimulus-evoked activity in correct auditory trials in W+ (green) and W- (magenta) contexts of selected target areas. (**F**) Same as in E, but for whisker trials. (**G**) Left: correlation between peak ΔF/F_0_ response to whisker and auditory trials in W+ and W- contexts. Right: correlation between peak ΔF/F_0_ responses in W- and W+ contexts for whisker and auditory trials. Each dot represents a mouse, dashed dark line represents equality of the two variables. (**H**) Left: Heatmap showing the first time at which the d’ comparing W- and W+ contexts in correct whisker trials passed above 2 for each pixel. Right: Same comparison, but derived from ROI time-courses shown in F.

To identify the earliest cortical area exhibiting context selectivity in whisker trials, we determined the time point at which each region reached a d′ greater than 2 between W+ and W− contexts. This analysis, performed both on pixel-wise activity maps and on the average fluorescence of target regions, revealed RSC as the first area to show significant context discriminability at 50 ms following the whisker deflection (Figure 3H). This rapid divergence of whisker stimulus evoked activity in RSC was observed consistently across individual sessions and mice (Figure 3 – figure supplement 3). To investigate whether differential movement related activity could contribute to this context divergence we analyzed the jaw and whisker movements in the first 200 ms after sensory stimulus for correct whisker and auditory trials in W- and W+ contexts (Figure 3 – figure supplement 4). This showed similar time courses for the whisker angle following sensory stimulus across contexts and a slightly higher whisker speed after whisker stimulus in the W+ context compared to the W- context, with d’ > 1 at times greater than 100 ms after the whisker stimulus. Therefore, differences in orofacial or whisker movements are unlikely to explain the observed effects of context on whisker-evoked neural activity.

Since the behavioral results suggested a progressive adaption following context transition (Figure 1F), we further investigated the changes of cortical activity at the context block transition time and over trials within each context block (Figure 3 – figure supplement 5). Focusing on a short timescale around context transition, we observed a rapid activation of auditory cortex within 100 ms, accompanied by a slightly stronger activation of RSC when transitioning from W+ to W- contexts than from W- to W+ contexts (Figure 3 – figure supplement 5A). To characterize the adaptation to context across trials following the context transition, we repeated the time course analysis (Figure 3F) including all whisker trials and taking into account the index in the block of each whisker trial (Figure 3 – figure supplement 5B&C). While stimulus evoked activity was relatively constant over trial number within each context block in wS1 and wS2, we observed a progressive decrease of RSC responses (as well as frontal cortex responses) to the whisker stimulus over trials in the W- context closely matching the behavioral results.

Together, our results show a context-dependent modulation of activity across the dorsal cortex before the onset of movement. They also point to RSC as a key region in relaying contextual information to other task-relevant cortical regions. The RSC is one of the main cortical outputs from the hippocampal formation (Li et al., 2025; Oh et al., 2014; Todd et al., 2019), which positions it as an interesting candidate to transmit contextual information to downstream cortical regions. To examine axonal projections, we injected AAV vectors into RSC using the coordinates defined in our optogenetic and imaging experiments, finding strong direct connectivity between RSC and frontal motor regions such as secondary motor cortex (wM1/2), consistent with previous anatomical studies (Oh et al., 2014; Yamawaki et al., 2016) (Figure 3 – figure supplement 6).

### Inter-area functional connectivity is modulated by context and upcoming choice

A possible mechanism for the same whisker stimulus to lead to licking or not depending on context could be to alter the functional connectivity between regions involved in the task. We hypothesized that this could be reflected in the correlation of the spontaneous activity during the pre-stimulus baseline period. To test this hypothesis, we computed seed-based correlations between each target area from the grid and each pixel from the image for each 2-s pre-stimulus window. For each seed area, we obtained a correlation map in both W+ and W- contexts for correct and incorrect trials (Figure 4A and Figure 4 – figure supplement 1). To identify significantly correlated pixels, we compared these correlation maps to null distributions obtained by shuffling entire blocks of trials and thereby preserving task structure (Harris, 2021). Finally, we reduced the obtained correlation map into the grid space which allowed us to build a graph connecting significantly correlated seeds in W- and W+ contexts for correct and incorrect trials (Figure 4B and Figure 4 – figure supplement 1A&B). We found a pattern of significant connectivity between whisker sensory regions, tongue-jaw orofacial regions and RSC with wM1/2 acting as intermediary node in the spontaneous pre-stimulus activity. We observed higher correlation values among orofacial regions, and between whisker motor and orofacial regions in W- compared to W+ contexts in the baseline preceding correct trials. Similarly, there was also an increase in correlation between wS1 and wS2 and between wS2 and wM1/2 in the W- context. Such contextual modulation of correlation was absent in baseline periods preceding an incorrect trial. This was due to an increase in correlation in incorrect trials in the W+ context (Figure 4B and Figure 4 – figure supplement 1C).

**Figure 4.**
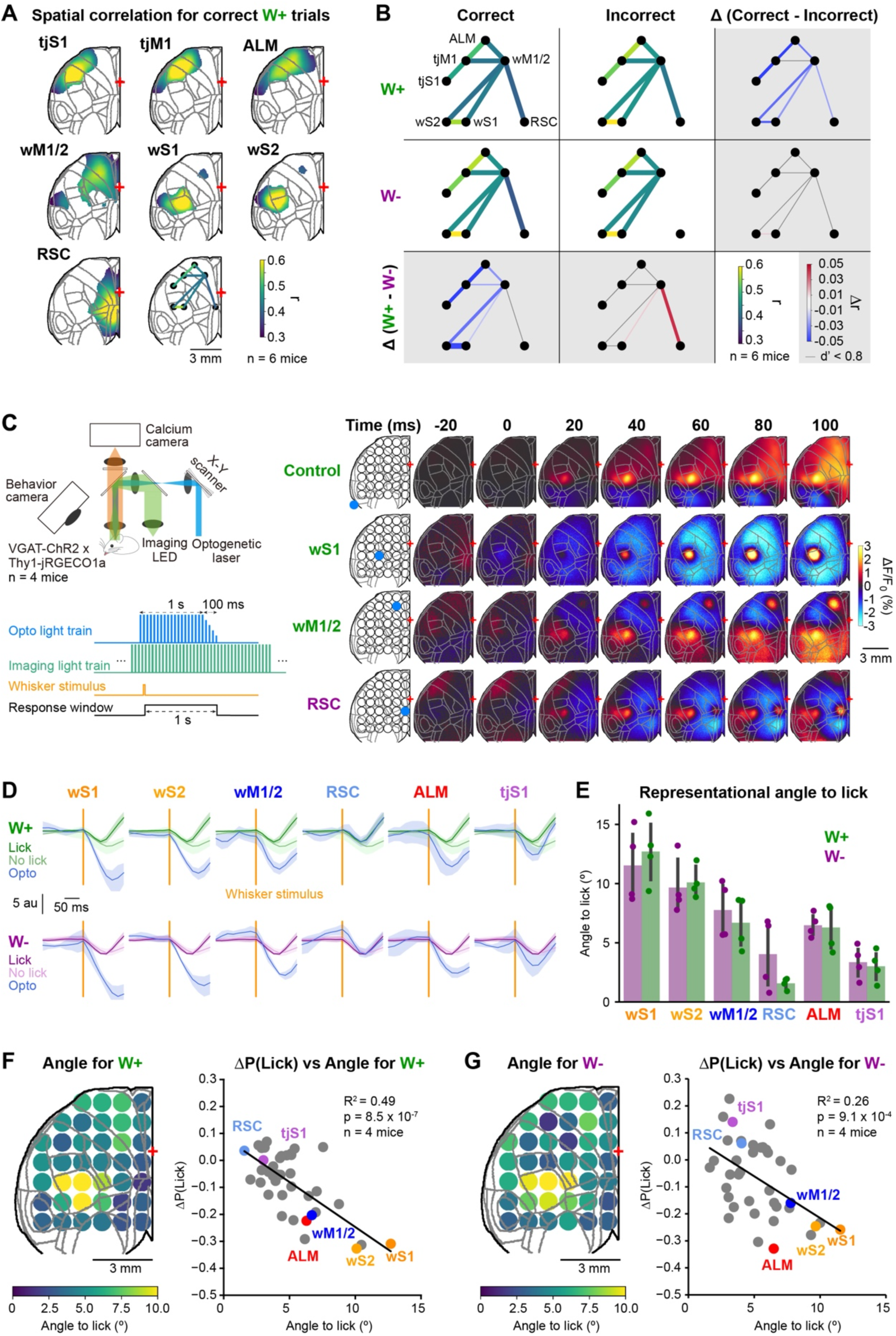
Context-dependent interactions across cortical areas. (**A**) Map of correlation values for pixels significantly correlated to each seed. (**B**) Graph of connected network for correct and incorrect trials in W- and W+ contexts extracted from maps of significant pixel to seed correlation. The rightmost column shows the difference between correct and incorrect trials in both contexts. The bottom row shows the difference between W+ and W- contexts for correct and incorrect trials. (**C**) Left: schematic of the experimental setup for simultaneous optogenetic and imaging. Right: averaged image series for whisker trials in W+ contexts for control trials and 3 example inhibition locations (blue dot on the grid). Time of frames in ms is aligned to whisker onset, and the optogenetic train starts in the immediately prior inter-frame interval. (**D**) Time course of activity projected on PC3 for whisker trials with optogenetic inhibition in W+ and W- contexts. Dark green lines show whisker trials with lick in the W+ context for control trials, whereas light green lines show the same for no–lick outcome. Dark purple and light purple lines show the equivalent in the W- context. Blue lines show the projected activity for optogenetic inhibition of each of the selected grid locations in W+ and W-contexts. Shaded areas represent 95% confidence intervals. (**E**) Quantification of the angle in PC3 between the optogenetic low dimensional projection and control-lick projection. Each dot represents a mouse and error bars are 95% confidence intervals. (**F**) Left: Grid projection of the average angle for each mouse between projections after optogenetic stimulation and control-lick projection in the W+ context. Right: Pearson correlation between the angle in the average map (see left panel) and changes in lick probability for the opto-widefield mice in the W+ context. (**G**) Same as F, but for the W-context.

Overall, this resting-state functional connectivity analysis showed reduced correlations between key cortical nodes in baseline periods preceding correct trials in the W+ context. Decorrelated cortical activity patterns can enhance information transmission through increased signal-to-noise ratios (Mukherjee et al., 2021; Poulet and Petersen, 2008; Shahsavarani et al., 2023), and the observed changes in functional connectivity could thus help propagate activity from whisker sensory cortex to frontal executive motor regions in the W+ context. It is important to note that these context-dependent changes in resting-state functional connectivity could relate to the overt context-dependent movements in the prestimulus baseline (Figure 1I&J) and/or a manifestation of higher-level internal rule representations.

### Optogenetic inhibition drives cortical activity modes predictive of behavior

So far, we have observed that inhibition of RSC prevents switching of the behavioral response to the whisker stimulus in the W- context. Importantly, inhibition of RSC in catch or auditory trials did not lead to any changes in performance, indicating that the presence of the context-sensitive stimulus is necessary for the effect of RSC inactivation to become apparent. We have also demonstrated that RSC is the first cortical region to display contextual differentiation in neural activity. Therefore, we investigated whether the behavioral effect of RSC inhibition could be explained by the optogenetically-driven changes in the neural activity patterns after whisker stimulus. To do so we combined calcium imaging of jRGECO1a and optogenetic inactivation in VGAT-ChR2 double-transgenic mice. We observed a diversity of cortical effects after inhibition during whisker trials (Figure 4C), which were not found in control mice only expressing jRGECO1a (Figure 4 – figure supplement 2A). For instance, inhibition of wS1 (which leads to a strong decrease in lick probability) resulted in widespread decrease in cortical activity, whereas inhibiting wM1/2, which also generated strong decreases in lick probability, concentrated the downstream inhibitory effect among motor areas while leaving the whisker sensory responses intact. These observations indicate a causal directionality of information flow, in which inhibiting each step in the sensorimotor transformation chain propagates selectively forward in the hierarchy. However, inactivation of RSC, which increases lick probability in whisker trials, did not increase activity in wM1/2 or tongue-jaw motor regions, suggesting more complex causal network dynamics. In order to shed light into the underlying patterns of neural activity supporting our observations, we reduced our cortical images into grid space and used PCA to define a low-dimensional space using the mouse averaged control whisker trials, separating lick trials from no-lick trials for each context to preserve the lick dimension (Figure 4 – figure supplement 2B-D). The loadings of the first principal components were uniformly distributed and could reflect a late movement driven activation distributed across all cortical areas (Figure 4 – figure supplement 2C&D). PC2 loadings show variation along the antero-posterior axis that could reflect differences between sensory and motor regions but its time course does not separate between lick and no lick in control conditions (Figure 4 – figure supplement 2C&D). The loadings of PC3 highlighted task-related cortical regions and its time course exhibited clear differences comparing lick and no-lick trials. We therefore projected trials with optogenetic inhibition of different cortical areas into this dimension to compare divergence from the control lick and no-lick neural trajectories (Figures 4D&E). For regions of interest that decreased task performance (wS1, wS2, wM1/2 and ALM), inactivation strongly deviated the trajectories away from the control-lick trajectory, towards no-lick. Only inhibition of RSC and tjS1, regions which led to increased licking, had trajectories that followed that of the control-lick in both contexts. To measure similarity of the opto-evoked neural trajectories with the control-lick trajectory, we computed the angle between these two traces. RSC and tjS1 – areas that increased licking in the W- context – had smaller angles with respect to the lick trajectories compared to the other regions, which decreased lick probability in all contexts (Figure 4E). This relationship held generally true across all inactivated regions in W+ and W- contexts (Figures 4F&G).

In summary, through combined calcium imaging and optogenetic inactivation, we found causal evidence for directional propagation of sensory signals to frontal cortex and revealed that RSC drives enhanced licking in W- contexts reflected in specific network subspace dynamics.

## Discussion

We developed a behavioral task where mice were able to switch their licking response to identical deflections of the C2 whisker based on an explicitly-presented auditory context (Figure 1). Unbiased optogenetic inactivation mapping of dorsal cortex confirmed key contributions of previously identified regions involved in whisker detection tasks (Chang et al., 2024; Esmaeili et al., 2021; Kwon et al., 2016; Le Merre et al., 2018; Oryshchuk et al., 2024; Sachidhanandam et al., 2013; Yang et al., 2016), as well as revealing an unanticipated role of RSC (Figure 2). Functional imaging of cortical activity with two different genetically-encoded calcium indicators each showed similar spatiotemporal dynamics of whisker sensory processing with the earliest context-dependent divergence in signalling being detected in RSC, out of the imaged dorsal cortical areas (Figure 3). Finally, functional connectivity measures and calcium imaging during optogenetic inactivation demonstrated causal contributions of RSC influencing the processing of whisker sensory information (Figure 4). Our results therefore point to an unexpected important contribution of RSC in our context-dependent whisker detection task.

While RSC has traditionally been linked to navigation (Mao et al., 2018, 2017) and vision (Kira et al., 2023; Sit and Goard, 2023; Vedder et al., 2017), recent work has indicated that RSC could also be important for tracking of reward (Hattori and Komiyama, 2022; Sun et al., 2021) and reward locations (Franco and Goard, 2021; Sun et al., 2021; Vedder et al., 2017). RSC is also important for the retrieval of contextual fear memory (Corcoran et al., 2011; Keene and Bucci, 2008; Robinson et al., 2012) and the optogenetic activation of memory-tagged RSC neurons was sufficient to reactivate fear memory (Cowansage et al., 2014). A growing body of evidence suggests that RSC is also involved in learning sensory associations for discrete cues (Todd et al., 2019). RSC was shown to contribute to negative patterning in behavioral tasks requiring rats to learn that the simultaneous presentation of two stimuli lead to an opposite outcome than each individual stimulus (Castiello et al., 2021). Finally RSC circuits are required for associating contextual information with appropriate motor output in a non-navigation task (Franco and Goard, 2021). Taken together with our observations, RSC appears as a key region for the integration of incoming sensory inputs to flexibly adapt a behavioral response according to the ongoing context.

In the future, it will be of great interest to investigate how cell-class-specific neuronal activity in RSC might contribute to orchestrating context-dependent interareal communication. Many important questions remain. The circuits signalling whisker sensory information and auditory contextual signals to RSC are currently unknown. Whisker sensory information could be relayed from wS1/2, with anatomical investigations suggesting that some L5IT cells in wS1 and wS2 directly innervate RSC (Liu et al., 2024). Context representations in hippocampus could be relayed to RSC through the direct CA1->RSC pathway (Cenquizca and Swanson, 2007), as well as indirect pathways via subiculum and entorhinal cortex (Sugar et al., 2011). These pathways could lead to the integration of whisker sensation with context in RSC, giving rise to enhanced excitation of a specific relevant subset of neurons in the whisker-rewarded context compared to the whisker-non-rewarded context.

An equally important question is how the activity of RSC neurons enables the context-dependent execution of the whisker-dependent detection task. First, further inactivation experiments would shed light on the timing at which RSC activity is necessary for the integration of contextual information. Specifically, it would be of great interest to inactivate RSC at different time points such as during the intertrial interval or at the transition between contexts. Second, because neurons in RSC project to many downstream brain regions, including wM1/2 (Figure 3 – figure supplement 6), it will be interesting to test the hypothesis that neurons with cell bodies located in RSC and with axons projecting to wM1/2 play a specific role in enabling context-dependent processing of the whisker stimulus. Key experiments will include: i) to measure the activity of individual neurons in RSC retrogradely-labelled from wM1/2 and/or image axons in wM1/2 originating from neurons in RSC; ii) to inhibit RSC->wM1/2 signalling via inhibitory opsins expressed in retrogradely-labelled neurons in RSC, for example with stGtACR2 (Mahn et al., 2018), and/or optogenetic presynaptic inhibition of RSC axons in wM1/2, for example using PdCO (Wietek et al., 2024); and iii) to stimulate RSC neurons projecting to wM1/2 to examine whether the auditory context can be substituted by optogenetic stimulation of specific subsets of RSC neurons. Finally, it is of course important to note that many subcortical regions (as well as non-dorsal cortical regions, which were not imaged) are likely to contribute importantly to context-dependent task performance.

## Methods

### Animals

All procedures were approved by the Swiss Federal Veterinary Office (License number VD-3769) and were conducted in accordance with the Swiss guidelines for the use of research animals. We used both male and female adult mice of at least 5 weeks at the time of surgery. For behavior only and pharmacological inactivation experiments we used wild-type mice. For calcium imaging experiments we used two subsets of mice: Thy1-jRGECO1a mice [Tg(Thy1-jRGECO1a)GP8.20Dkim/J, JAX: 030525] (Dana et al., 2018), which express a red shifted calcium indicator broadly across cortical regions, and Rasgrf2-dCre mice [B6;129S-Rasgrf2 <tm1(cre/folA)Hze>/J, JAX: 022864] (Harris et al., 2014) crossed with Ai148 mice [B6;Ai148(TIT2L-GC6f-ICL-tTA2)-D, JAX: 030328] (Daigle et al., 2018), which express the green fluorescent calcium indicator GCaMP6f specifically in excitatory layer 2/3 neurons. To test the effect of optogenetic stimulation of inhibitory neurons, we used VGAT-ChR2 mice [B6.Cg-Tg(Slc32a1-COP4*H134R/EYFP)8Gfng/J, JAX: 014548] (Zhao et al., 2011).

For simultaneous calcium imaging and optogenetic manipulations we crossed VGAT-ChR2 mice with Thy1-jRGECO1a mice. For GCaMP calcium imaging control experiments, we used GAD67-GFP mice [CR.Cg-Gad1tm1.1Tama/Rbrc, MGI:3590301] (Tamamaki et al., 2003), which have a similar level of green fluorescence. For jRGECO1a calcium imaging control experiments, we used PV-Cre mice [B6;129P2-Pvalbtm1(cre)Arbr/J, JAX: 008069] (Hippenmeyer et al., 2005) crossed with R26_LSL_tdTomato mice [B6.Cg-Gt(ROSA)26Sor<tm9(CAG-tdTomato)Hze>/J, JAX: 007909] (Madisen et al., 2010), which have a similar level of red fluorescence.

### Implantation of headpost and skull preparation

Mice were first implanted with a metal head-post under isoflurane anesthesia (1.5-2.5 % isoflurane). Before the start of the surgery, we injected carprofen (7.5 mg/kg at 1.5 mg/ml, subcutaneous) for general analgesia, and a mix of lidocaine and bupivacaine below the scalp as local analgesic. An ocular ointment (Vita-Pos, Pharma Medica AG, Switzerland) applied over the eyes prevented them from drying during the surgery, and a solution of povidone-iodine (Betadine, Mundipharma Medical Company, Bermuda) was applied for skin disinfection before incision. We monitored body temperature throughout the surgery and it was maintained at 37°C with a heating pad (ThermoStar, Intellibio, France). After excising the scalp with surgical scissors to expose the skull, we thoroughly cleaned it with cotton buds and a scalpel blade to remove the periosteal tissue. After further disinfection with Betadine, the skull was dried with cotton buds and made optically clear using a thin layer of super glue applied over the dorsal part of the skull (Loctite super glue 401, Henkel, Germany). We then glued a custom-made head fixation implant above the right hemisphere, and further secured it with self-curing denture acrylic (Paladur, Kulzer, Germany or Ortho-Jet, Lang, USA). Finally, a second thin layer of the glue was applied homogeneously on the left hemisphere to ensure a smooth and equally transparent field of view. Particular care was taken to ensure that the left hemisphere of the dorsal cortex was free of denture acrylic and only covered by super glue for optical access. This intact, transparent skull preparation was used to perform intrinsic optical signal (IOS) imaging, widefield imaging experiments, and opto-inhibition screening. Mice were returned to their home cages and ibuprofen (Algifor Dolo Junior, Verfora SA, Switzerland) was added to the drinking water for three days after surgery.

### Behavioral task and training curriculum

We trained head-fixed, water-restricted mice to perform a context-dependent multisensory detection task. During the behavioral experiments, all whiskers were trimmed except for the C2 whisker, and the mice were water restricted to 0.9-1.2 mL of water/day (depending on the initial body weight). Mice were trained daily with one session per day, their weight and general health status were carefully monitored using score sheets. Mice were habituated to be head-fixed to a metal post with their head held in a 30-degree angle. During the first session, mice were also habituated to licking from a water spout positioned on their right side which delivered 5 µL of water in response to some licks. Next, we trained them in an auditory detection task for three to five days during which licking in a 1 s response window after the auditory cue (10 ms, 10 kHz tone of 74 dB embedded in the continuous background white noise of 80 dB) was rewarded. Inter-trial interval was uniformly sampled between 6 and 10 s including a final 2 to 5 s no-lick window during which the mice were required not to lick for the trial to be initiated. Once the mice showed clear detection of the auditory cue, we introduced for two to three days a C2 whisker deflection (a 25 mT magnetic pulse driven by a 3 ms cosine pulse sent into an electromagnetic coil positioned under the head of the mouse and acting upon a small metal particle attached to the C2 whisker), which like the auditory cue was also rewarded if the mice licked within 1 s following the stimulus. Mice were then trained in a block-based contextual task. Contextual information was provided by alternating pink and brown noise instead of white noise in the background. Each mouse was randomly assigned a whisker-rewarded (W+) context at the beginning of training. Pink and brown background noise alternated every block of 20 trials, and was continuously present and available to the mice. Within each 20-trial block, mice received 8 whisker trials, 4 auditory trials and 8 catch trials presented pseudo-randomly. Mice were trained in this paradigm until they reached expert level. To determine expert level, we computed a contrast value for each block of context apart from first and last ones. It was defined as the average of the absolute difference in the whisker hit rate between one block and the two surrounding blocks. For each session we obtained a distribution of contrast values and compared the mean to a threshold of 0.375, which essentially requires mice to respond differently for at least 7/16 possible whisker trials between blocks throughout the entire session. Sessions with a lower bound 95% confidence interval of mean contrast values (estimated from bootstrapping 10,000 times) above the 0.375 threshold were classified as expert. For each session we computed the context discriminability using Cohen’s d.

### Video tracking and pose estimation

Simultaneously with behavioral training and imaging, we recorded mouse orofacial movements using two cameras (DMK 37BUX273, The Imaging Source) positioned above and on the side of the mouse face with pixel size calibrated to real distance (top camera 24 pixels/mm, side camera 28 pixels/mm). Acquisition was freely running at either 100 Hz or 200 Hz and we recorded timestamps of each image relative to session start-time. We trained a network using the behavioral pose estimation software DeepLabCut (Mathis et al., 2018) (v3.0.0rc6) to track different body parts of the orofacial region. We labelled 940 frames uniformly extracted from 47 sessions in 24 mice for each camera view. We trained our network using 95% of the data as training fraction, and a ResNet 50 with a batch size of 8 (default parameters). After training, we obtained a train error of 3 pixels at 60% likelihood cut-off, and a test error of 3.71 pixels for the same cut-off value in the side view camera. Evaluation results were similar for the top view camera, with 3.8 pixels of error in the train dataset, and 3.75 pixels for the test dataset at 60% likelihood cut-off.

### Optogenetic inactivation

For optogenetic inactivation experiments, a multimode custom patch fiber (0.22 NA, 105 μm diameter core, FG105UCA, Thorlabs, USA) attached to a 473 nm laser source (S1FC473MM, Thorlabs, USA) was used to direct a laser beam into a set of galvo-galvo scan mirrors (GVS202, Thorlabs, USA). We designed a grid of 39 points separated by 1 mm covering the left dorsal cortex, and calibrated the grid to bregma. Due to the large number of stimulation points, we further randomly divided the total grid into three subgrids, which each mouse experienced over multiple sessions in a randomized rotating schedule. In fifty percent of the trials, we randomly directed the laser beam to a subgrid point to excite ChR2 expressing neurons with a train of 5 ms square pulses at 50 Hz from 10 ms before the stimulus to 1 sec after. The other fifty percent of trials were non-stimulated trials, in which we directed the laser beam outside the brain. We iterated over subgrid points over a single session and over subgrids over multiple sessions (days).

### Pharmacological inactivation

We performed muscimol inactivation on mice trained in the context-dependent detection task after they reached expert level for at least two consecutive days. On the day before injection, after the training session, we performed a small craniotomy above the targeted region. We used the intrinsic optical signal to target the C2 column of the whisker primary sensory cortex (C2-wS1), the forepaw primary sensory cortex coordinates were estimated 1.5 mm anterior and 1.5 mm medial compared to C2-wS1, the retrosplenial cortex coordinates were 1.5 mm posterior to bregma and 0.5 mm lateral from the midline. On the inactivation day, we injected 100 nL per injection depth of muscimol 5 mM diluted in Ringer (or Ringer alone in control experiments) at 200, 400, 600 and 800 µm below the pia. We started the behavioral session fifteen minutes after the last injection. We performed up to four injections (muscimol and/or control) per mouse separated by at least one day with a normal behavioral session.

### Calcium imaging

We collected cortex wide imaging data from 4 mouse lines expressing different fluorescent proteins: i) mice expressing the red calcium indicator jRGECO1a; ii) mice expressing tdTomato as a calcium insensitive red fluorophore control for jRGECO1a; iii) mice expressing the green calcium indicator GCaMP6f in layer 2/3 excitatory neurons; and iv) mice expressing GFP as a calcium insensitive green fluorophore control for GCaMP6f. We excited jRGECO1a (and tdTomato) using 563 nm light (19.2 μW/mm^2^ on the cortical surface; 567 nm LED, SP-01-L1, Luxeon, Canada; 563/9 nm BrightLine HC bandpass filter, Semrock, USA). Red emission light was separated from excitation light using a dichroic mirror (Beamsplitter T 588 LPXR, Chroma, USA) and bandpass filtered (645/110 ET Bandpass, Semrock). To excite GCaMP6f (and GFP), we directed blue light from a 488 nm LED (M490L4, Thorlabs), narrowband filtered at 488 (488/6 BrightLine HC, Semrock, USA). Emitted light was separated from excitation light using a beamsplitter (T 495 LPXR, Chroma, USA) and bandpass filtered (525/50 BrightLine HC, Semrock, USA). Images were taken using a 16-bit monochromatic scientific Complementary Metal Oxide Semiconductor camera (sCMOS, Hamamatsu Orca Flash 4.0 v3, Hamamatsu Photonics, Japan) coupled in face-to-face tandem to an objective and imaging lens (Nikkor 50 mm f/1.2, Nikon, Japan; 50 mm video lens, Navitar, USA, respectively). We collected 256 x 320 pixel-sized images of the dorsal cortex of the brain at 100 Hz with a 10 ms exposure time using the “SynchronousTrigger” mode.

For combined calcium imaging and optogenetics, we collected 256 x 320 pixel-sized images of the dorsal cortex of the brain at 50 Hz with a 2.5 ms exposure time in the “EdgeTrigger” mode. Imaging frames were interleaved to the optogenetic pulses to minimize optolight contamination.

### Axonal projections from retrosplenial cortex

To visualise axonal projections from the retrosplenial cortex we injected an adenoassociated virus (AAV) allowing for the expression of GFP fused to the chronos opsin to favor axonal labelling (AAV5.Syn.Chronos-GFP.WPRE.bGH, titer: 3.82 x 1013 vg/mL, AV-5-PV3446, University of Pennsylvania Vector Core). Virus injections were performed 0.5 mm lateral to the sagittal sinus and 1.5 mm posterior to bregma. AAV was injected at 800, 600, 400, and 200 µm below the pia (25 nL of 1:10 dilution of stock virus per injection site for a total volume of 100 nL). Mice were perfused with 4% PFA 6 weeks after the injection and the brains were cleared following the iDISCO protocol (Liu et al., 2024; Renier et al., 2014). Brains were dehydrated (methanol/dH_2_O gradient), delipidated (dichloromethane), bleached (hydrogen peroxide) and then permeabilized and blocked (bovine serum albumin) in preparation for immunostaining. Samples were incubated in a primary antibody solution (Rabbit anti-GFP Ab290, Abcam, 1:1000 dilution) for 7 days and after several washes incubated for a week in a secondary antibody solution (Alpaca anti-Rabbit Alexa 647, SA5-10327 ThermoFisher, 1:800 dilution). After a final step of dehydration and delipidation, samples were immersed in ethyl cinnamate for refractive index matching. Dual side light sheet imaging was performed with a Zeiss Z1 microscope with a resulting voxel size of 1.58 x 1.58 x 5.34 µm, where the xy plane corresponds to horizontal sections. Finally, samples were registered to a mouse brain atlas optimized for light sheet fluorescence microscopy (Perens et al., 2021).

### Quantification and data analysis

#### Computation of ΔF/F_0_

To compute relative changes in fluorescence over time, for each pixel in the widefield video, we computed the 5^th^ percentile of the distribution of session-wide values and used that as F_0_. We then computed the moment-by-moment difference between the fluorescent values and F_0_ and divided by F_0_. In Figures 3&4 and Figures 3&4 supplement 1, calcium data was further processed by subtracting the average of the 250 ms prior to stimulus onset to the stimulus aligned images for each trial before averaging.

### Behavioral kinematics

After obtaining the session-wide pose estimation from DeepLabCut, we computed body part kinematics prior to integration in NWB datafiles.

We computed the opening of the jaw as the vertical distance from a reference point determined by the 5^th^ percentile of the entire jaw trace (after filtering at 50% likelihood cut-off). Jaw speed was then computed as the absolute of the first derivative of the jaw opening traces.

To compute the whisker angle, we calculated the arc cosine of the dot product between the vector formed by the metal particle used to deliver the whisker deflections and the base of the whisker, with the vector crossing the midline of the mouse (computed by the median position over the session of the tip and base of the nose) divided by the norm:

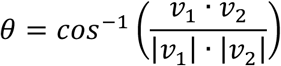

where v1 = (X_particle_ – X_whisker_base_, Y_particle_ – Y_whisker_base_), and v2 = (median(X_nose tip_) – median(X_nose_base_), median(Y_nose_tip_) – median(Y_nose_base_)). Finally, this angle was rotated such that a fully protracted whisker (parallel to the body) would be 180 degrees and a fully perpendicular whisker to the midline was 90 degrees. As with the speed of jaw movements, whisker speed was computed as the absolute of the first derivative of the whisker angle.

In addition, we computed the area of the pupil as the surface of the polygon with the four points tracking the top, right, bottom, and left of the pupil as vertices. In order to be computed, all four points were required to have a likelihood bigger than 0.5 for each given point.

### Seed correlations

After extracting the minimal common quiet window period before stimulus onset from each trial (2 s quiet window), we progressively iterated over regions of interest (wS1, S2, wM1, RSC, ALM, tjM1, and tjS1) and pooled the pixels inside the 1 mm circle centred on the coordinates of the optogenetic grid inhibition. As such we obtained a source matrix for each ROI being (trials x frames), and a target tensor for the entire session being (trials x frames x pixels). Then we iterated over pixels to compute the trial-by-trial Pearson’s zero lag correlation between the ROI and each pixel.

To generate a null distribution while preserving general structure of the session, we shuffled 1,000 times entire blocks by randomly swapping the block identity, which preserved the within-block structure. This shuffling method is stricter than simply shuffling all trials, since it would preserve slow components of the block. We considered pixels significantly correlated to the seed if the correlation coefficient of that pixel was on average larger or equal than 1.8 times the standard deviation of the null distribution.

### Principal component analysis

To compute the low dimensional space onto which to project our simultaneous widefield imaging and optogenetic inactivation data, we first extracted the widefield images for the time period of 300 ms surrounding a whisker trial and reduced them to grid space by pooling the pixels within a 1 mm diameter circle centred on locations from our optogenetic grid. We then extracted control stimulation trials (where light is projected outside the brain, ∼50% of total trials), and computed the total average ΔF/F_0_ by first averaging across each individual mouse pseudosession and then across mice. Next, we computed the principal components of the matrix that has as columns the optogrid ROI ΔF/F_0_, and as rows a multi-index of time, lick, trial type (whisker or catch), and context. Preserving the lick dimension allows us to investigate the differences of lick trials vs no-lick trials. We then recomputed the mouse average matrix by averaging by trial type within each pseudosession regardless of lick or no lick (thus computing the weighted average calcium responses best mimicking the effect of our inhibition in Figure 2). Then, we projected the mouse average data using the same set of weights obtained by projecting the control trials.

### Gradient boosted decision tree model

Behavioral data from all expert sessions were concatenated in a single table. One single session was randomly chosen to be used as a held-out test session. Then, data were split into a 67% train set and 33% test set, with a stratification strategy to ensure that the proportion of the binary outcome was the same in both train and test sets. The target value was whether the animal licked or not in response to the whisker stimulus. A Gradient Boosted Tree (GBT) model was fitted on the data using the package XGBoost (Chen and Guestrin, 2016).

Gradient Boosted Trees are ensemble learning methods that build models sequentially, where each tree is a weak decision tree classifier trained to predict the outcome of a trial. The key principle is that each subsequent tree is trained to correct the prediction errors made by the previous trees, with the final prediction being the weighted sum of all tree predictions. This iterative approach allows the model to progressively improve its performance by focusing on the most challenging cases. GBT models have been demonstrated to be particularly effective for tabular data, often outperforming nonlinear neural network-based classifiers (Grinsztajn et al., 2022). Furthermore, this approach has proven valuable for modeling behavioral data as it can capture non-linear relationships as well as complex interactions that traditional linear models fail to capture (Griffiths et al., 2024).

The model was trained on 14 features spanning two categories. First, task-related features included the mouse identity, the context (W+ or W−), cumulative water received, whether the previous trial was correct (hit or correct rejection), whether the previous trial was rewarded (i.e., a hit), whether the previous whisker trial was rewarded, the type of the last stimulus (auditory or whisker), the number of whisker trials elapsed in the current context block, and the time elapsed since the last auditory or whisker stimulus (excluding catch trials). Second, movement features were derived from the kinematic variables described above (pupil area, Jaw y, Jaw speed, whisker angle, and whisker speed), each summarized as their mean value over the 2-second window preceding stimulus onset.

Hyperparameter tuning was performed using Bayesian optimization implemented in Optuna (Akiba et al., 2019). The optimization minimized binary log loss for classification tasks using a holdout validation approach, where the training data was split 80/20 into training and validation sets. Early stopping with 100 rounds was applied to prevent overfitting during model training. The search space included 12 hyperparameters. The best hyperparameters were then used to train the GBT model on the full training set, using binary log loss. Balanced accuracy was then computed on the test set. To interpret the trained GBT model and gain a better understanding of how each feature contributes to the prediction, we used SHapley Additive exPlanations (SHAP) analysis (Lundberg and Lee, 2017). In short, SHAP values, originating from cooperative game theory, provide a mathematically rigorous method for attributing the contribution of each feature to a model’s prediction.

### Statistics

For statistics we used parametric tests and correction for multiple comparisons using the Bonferroni method.

### Figure-specific analysis methods

Figures 1C-E: These panels describe the average probability of licking to different trial types of expert mice in the context task. Each dot represents a mouse, obtained by first averaging trials within a session then across sessions for each mouse. Population average is obtained by further averaging over mice, and error bars show the 95% confidence interval of the population mean.

Figure 1F: This panel represents the speed of switching behavior after context transition. Each dot represents the population average lick probability as a function of the whisker trial indices, before and after context transition; error bars show the 95% confidence interval of the population mean.

Figure 1G: Summary plot representing the speed of switch in the last vs first trial after context switch. Each connected pair of dots represents a mouse at whisker trial -1 and +1 extracted from Figure 1F.

Figure 1H: Same as Figure 1F but represented as time around context transition in which the last/first whisker trial occurs. We selected for each session the first and last whisker trials of each block around context transition (i.e., excluding first whisker trial of the first block and last whisker trial of the last block), this yielded approximately 10 ‘first’ whisker trials and 10 ‘last’ whisker trials for each session. We then assigned a label to each of these trials according to which of the 10-s bins around context transition it belonged. We repeated the same process over all sessions. For each 10-s time bin we defined the lick probability as the number of trials with lick divided by the total number of trials in this time bin.

Figures 1I-J: Quantification of postural and movement changes according to context and lick. We extracted movement traces for jaw opening, whisker angle and pupil area from the NWB files. Pupil area was computed as the surface of the polygon that has as vertices 4 points surrounding the pupil (top, bottom, left and right), and only computed for timestamps in which the likelihood values of all vertices were above 50%. We computed whisker and jaw speed as the absolute of the derivative of the jaw and whisker traces respectively, times the frame rate. Next, we filtered the body parts using an adaptive threshold of 80% likelihood for whisker, 60% for jaw, and 50% for pupil and any body part that kept less than 70% of the data points after filtering over the entire session was discarded from analysis. We then extracted the two-second arrays corresponding to the pre-stimulus imposed quiet window of whisker trials and averaged them within sessions, then within mouse preserving context and lick categories. To visualize the changes in posture or movement, we subtracted for each mouse the average value of the No-lick, W- trials. We used the population array to compute statistics using paired t-tests. We computed paired effect size as the absolute difference between means divided by the standard deviation of the individual differences: 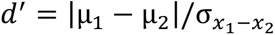, where µ signifies the mean and σ the standard deviation.

Figure 1K: For our Gradient Boosted Decision Tree model, we show the results of the permutation test with scikit-learn (Pedregosa et al., 2011) used to validate which features have the largest contributions to model performance. In this procedure, the feature importance was assessed by randomly shuffling each feature independently 10,000 times while keeping all other features intact. For each feature, the mean decrease in balanced accuracy of the test set relative to the unpermuted test set balanced accuracy was recorded across all permutations as a measure of that feature’s importance.

Figure 1L: In this panel, we show the interaction between the SHAP values of the feature “whisker trial in block” versus the position of the whisker trials in the block as it is used in Figure 1F. To compute the SHAP values, we used the Python SHAP package (Lundberg and Lee, 2017) with the interventional feature perturbation method. This method treats each feature’s contribution as if we were experimentally manipulating that feature while holding others constant, which is particularly important when features are correlated or interdependent. To achieve this, we randomly sampled 100 trials from the training data to serve as a background dataset representing the typical distribution of our data. For each prediction, the interventional method calculates SHAP values by replacing each feature’s value with values from the background dataset while keeping other features fixed. This process estimates the expected change in model output when intervening on a specific feature. The SHAP value for a feature in a given trial represents the difference between the model’s prediction with the actual feature value and the expected prediction when that feature is marginalized over the background distribution. Positive SHAP values indicate that a feature increases the probability of licking for that specific trial, while negative values indicate a decrease. The violin plots represent the distributions of SHAP values for each whisker trial in block, and context, from single trials included in the model.

Figures 2C-E: Summary plots representing the changes in lick probability with respect to control stimulation in our optogenetic manipulation experiments. Given that our protocol for optogenetic inactivation completes a grid over the course of ∼3 sessions, we considered all the sessions for each mouse as a single pseudosession. For each mouse, context, trial type and inactivation coordinates, we computed the probability of licking (see also Figure 2 – figure supplement 1A&B) and subtracted from it the probability of licking in our control stim condition (where light is projected outside the brain), thus obtaining for each grid coordinate a ΔP(Lick) for each mouse. To estimate the effect size of inactivation, we shuffled the labels of entire blocks of trials (keeping trial order and structure, but shuffling block identity) and recomputed ΔP(Lick) in the same way to obtain a null distribution. We then computed the d’ between the null distributions of ΔP(Lick) and the observed distribution.

Figure 2F: Cortical maps representing the changes in lick latency recorded from the piezo sensor attached to the water spout. We computed for each mouse the average of the spout contact time in response to auditory trials in both contexts and to whisker trials in the W+ context for each inhibition location and subtracted the averaged spout contact time from control trials. We further averaged this distribution of change in spout contact time over mice. This resulted in the grid maps shown in the top row. To provide an estimate of the variance we computed for each inhibition site the distance (d’) between the distribution of spout contact time to the one obtained in control trials. This resulted in the grid maps shown in the bottom row.

Figure 3B: Average jaw opening trace after whisker onset. As in Figures 1I&J, we extracted movement traces for jaw opening from the NWB files and filtered them at a 60% likelihood threshold. We extracted jaw opening traces centred at the time of whisker delivery for correct trials (i.e., licks to whisker in the W+ context and no licks in the W- context) and centred them by subtracting the average jaw opening value 250 ms before stimulus onset. We averaged the jaw traces first within session and then across sessions for each mouse before computing the population average response across mice. The shaded region represents 95% confidence intervals computed over mice.

Figure 3C: Average reaction times for each mouse computed from the jaw traces. For each auditory and whisker trial with lick in both W+ and W- contexts, we computed the standard deviation of the 2-second baseline window preceding the stimulus and detected the first time point going above three time this value following the sensory stimulus. We averaged reaction time over trials within individual session then across sessions for each mouse. Population average is obtained by further averaging over mice, and error bars show the 95% confidence interval of the population mean. Each dot represents a single mouse.

Figure 3D: To map sensory-evoked activity onto the grid of coordinates, we manually drew a 1 mm circular ROI centred on the peak response to whisker or auditory trials for each imaging session. We then averaged the centre of mass of wS1, wS2 and A1 first across sessions for each mouse and then across mice. This averaged center of mass is represented by small stars on the grid.

Figures 3E**&F:** Population average stimulus-evoked calcium dynamics across the dorsal cortex (left) and from selected regions of interest (right). We first realigned activity (normalized as ΔF/F_0_) to trial onset for each session and subtracted the baseline activity (defined by the average value of the 50 ms preceding the trial) in a trial-based manner. We then averaged this normalized activity over trials within each session. We further averaged over sessions for each mouse and finally over mice. The d’ rows show the distance for each pixel between the distributions of mouse averages for W+ and W- blocks (6 values for each context for each pixel). For the ROI time-courses, the error bars shadowing the population averages show the 95% confidence interval of the population mean.

Figure 3G: Representation of similarity of responses across contexts or trial types. We used the mouse averaged response to auditory and whisker trials in W- and W+ contexts to detect the maximal ΔF/F_0_ value in a time window from 50 ms to 120 ms after stimulus delivery. We show the correlation of maximal ΔF/F_0_ in this time window between the two sensory modalities in each context or between the two contexts for each sensory modality. Each dot represents a mouse, each color represents a target region.

Figure 3H: (Left) Map representing the latency to context divergence for each pixel, defined as the time of the first frame at which the d’ (shown in Figure 3F bottom row) exceeds a value of 2. Since d’ can be sensitive to low variance time points, the first 20 ms after the stimulus delivery - before any response in both contexts - were excluded from the search. (Right) Ranking of regions of interest showing the earliest contextual divergence: same as Left panel, but with d’ computed from the ROI averages shown on the right in Figures 3E&F.

Figures 4A**&B:** Summary results of the seed correlation analysis. For each imaging session, we extracted the ΔF/F_0_ widefield data in the 2-s quiet window preceding each trial start. We also extracted the same time periods for the pixel averaged ΔF/F_0_ traces corresponding to the area under a given optogenetically defined ROI to obtain our seeds and computed the zero-lag Pearson correlation coefficient between the seed and each pixel in the widefield image, for each trial independently. We focused our analysis to the seed coordinates that showed effects of inactivation in whisker trials (wS1, wS2, RSC, wM1/2, ALM, tjM1, and tjS1). We also shuffled entire blocks of trials (to preserve session structure) by shuffling the block id’s and recomputed the zero-lag Pearson correlation coefficient 1,000 times to obtain a null distribution. For each pixel, trial, and session we then computed the number of standard deviations away from the shuffle distribution that a given pixel correlation coefficient was, which allowed us to compute an estimate of significance. We then averaged first within session, then over sessions for each mouse the correlation maps (reconstructed into widefield cortical maps) maintaining the context, and correct vs incorrect categories. To construct connectivity maps, we masked the pixels that had on average, for each category, a correlation coefficient larger or equal to 1.8 times the standard deviation of the shuffle distribution, which under consideration of a gaussian distribution corresponds to the 95^th^ percentile. Then, we reduced the masked coefficient cortical map into optogenetic grid space by averaging the masked correlation coefficients of the pixels under the area corresponding to the 1 mm circle centred on optogenetic coordinates. Finally, we connected the graph by cross validating which nodes of the optogenetic coordinates were significantly correlated (>=1.8 times the standard deviation of the null distribution) for each seed. Once we had a connected graph for each context, and correct vs incorrect trial types, we subtracted the r values for each connected pair to obtain Δr maps. We also tested for significance of the difference by means of a related t-test corrected for multiple comparisons using Bonferroni correction, and computed the effect size of the difference by computing the Cohen’s d for related samples as explained above: 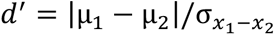. Each plot in Figures 4A&B corresponds to the average correlation r values across mice, and the line thickness in the Δr maps in Figure 4B increases with d’ such that thinnest colored lines correspond to a d’ of 0.8 and max thickness is capped at 2. If the correlation difference between two points has a d’ below 0.8 this line is plotted in gray.

Figure 4C: Description of the combined widefield and optogenetic setup and example image sequences. To obtain average post-optogenetic stimulation widefield time courses we averaged the trial start aligned widefield ΔF/F_0_ arrays for each session, context, and optogenetic coordinate point. Then we averaged over sessions for each mouse and finally across mice to obtain mouse average widefield maps.

Figure 4D and Figure 4 – figure supplement 1E-H: Results of the dimensionality reduction analysis on the combined widefield and optogenetics data. To compute low dimensional projections of our simultaneous optogenetic and widefield experiments we first separated the control trials (where light was directed outside the brain) from the optostim trials. We extracted the optogenetic grid space widefield images (obtained by averaging pixels inside the 1 mm circle centred to each coordinate in our optogenetic grid) aligned to trial onset in whisker and catch trials. We then computed the population average control trials by averaging first all the control trials in the pseudosession generated for each mouse, and then averaging across mice while preserving the context, trial type, lick, and timestamp categories, as well as the identity of each coordinate point. We standardized our dataset by fitting and then transforming using the StandardScaler from sklearn, and pivoted to obtain a matrix with the optogenetic grid coordinates as columns and context, trial type, lick, and timestamps as rows before computing the 15 first principal components of this matrix using PCA. In Figure 4 - figure supplement 1E&F, we plot the variance explained and the weights mapped back into optogenetic grid coordinates for the first 3 components. Once we obtained our principal components, we repeated the same process for the stim trials, but in this case, we preserved mouse identity and coordinate of the optogenetic stimulation, hence obtaining a matrix that had as columns the optogenetic grid coordinates, and mouse identity, optogenetic stimulation coordinate, context, trial type, and timestamp categories as rows. This way, our average projections will contain the weighted average over the lick axis for the stim conditions and will reflect better the observations in the average optogenetic inhibition results. We standardized the stim matrix using the same parameters to scale the control matrix, then projected onto the same low dimensional space using the control principal components. We then aggregated over mice for each context, optogenetic stimulation coordinates, and principal component to obtain the low dimensional projections for each PC as found in Figure 4D and Figure 4 - figure supplement 1H. Then, we plotted the control-lick and control-no-lick projections for each context, and the projection obtained from the optoinhibition trials, with shaded area representing 95% confidence interval computed over mice.

Figures 4E-G: Summary plots representing the difference between the low dimensional projections in control-lick versus photoinhibition. After obtaining the low dimensional projections for the control and stim mouse average trials, we computed the angle between the stim traces for each mouse and the average control-lick trace to estimate the divergence from lick. We computed the angle in radians by obtaining the cosine of the dot product between the control-lick and stim vectors, divided by the norms, and then convert it to degrees:

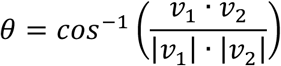

Figure 4E contains a bar graph representation of the angles for the selected subset of optogenetic grid coordinates that show effects in Figure 2. Each dot represents the angle between control-lick and optoinhibition low dimensional projection in PC3 for each context, and error bars represent 95% confidence intervals. Figures 4F&G(left) contain the mouse average angle between control-lick and optoinhibition PC3 projections for each context respectively for each optogenetic stimulation coordinate. Figures 4F&G(right) show the linear regression between the value of the angles in PC3 for each point and the evoked changes in lick probability for the subset of mice that we conducted the simultaneous optogenetic and widefield experiments.

## Data availability

For each imaging session, imaging data, behavioral data, cortical region contours and calcium traces, were combined into a single NWB file. NWB offers a common format for sharing and analyzing neurophysiology data (Rübel et al., 2022). Subsequently, we developed open-source Python scripts to analyze data in the NWB format. The full dataset in NWB format will be available on the DANDI archive and processed data will be available via Zenodo at https://doi.org/10.5281/zenodo.17424306.

## Code availability

Code for data acquisition and behavior control is available on github (behavior GUI repo). All the code used to preprocess and convert the data into NWB format is available on github (NWB converter repo), all the code used for data analysis is available on github (NWB analysis repo), Cicada is available on gitlab. The Python code used for analyses will also be available via Zenodo at https://doi.org/10.5281/zenodo.17424306.

## Acknowledgements

We thank the members of the Petersen laboratory for helpful discussions. This work was supported by grants from the Swiss National Science Foundation: 31003A_182010 (C.C.H.P.), 310030_219343 (S.C. and C.C.H.P.) and TMAG-3_209271 (C.C.H.P.).

## Author contributions

P.B., R.F.D., S.C. and C.C.H.P. conceptualized the study; P.B., R.F.D., A.B. and A.R. wrote the data acquisition codes. P.B. and R.F.D. performed all animal experiments and constructed the database. R.F.D. and L.S. performed the anatomy experiments. P.B., R.F.D., and J.L. analyzed the data. P.B., R.F.D., S.C. and C.C.H.P. wrote and edited the manuscript, with comments from all authors; S.C. and C.C.H.P. acquired funding and provided overall supervision.

---

**Figure 1 – figure supplement 1.**
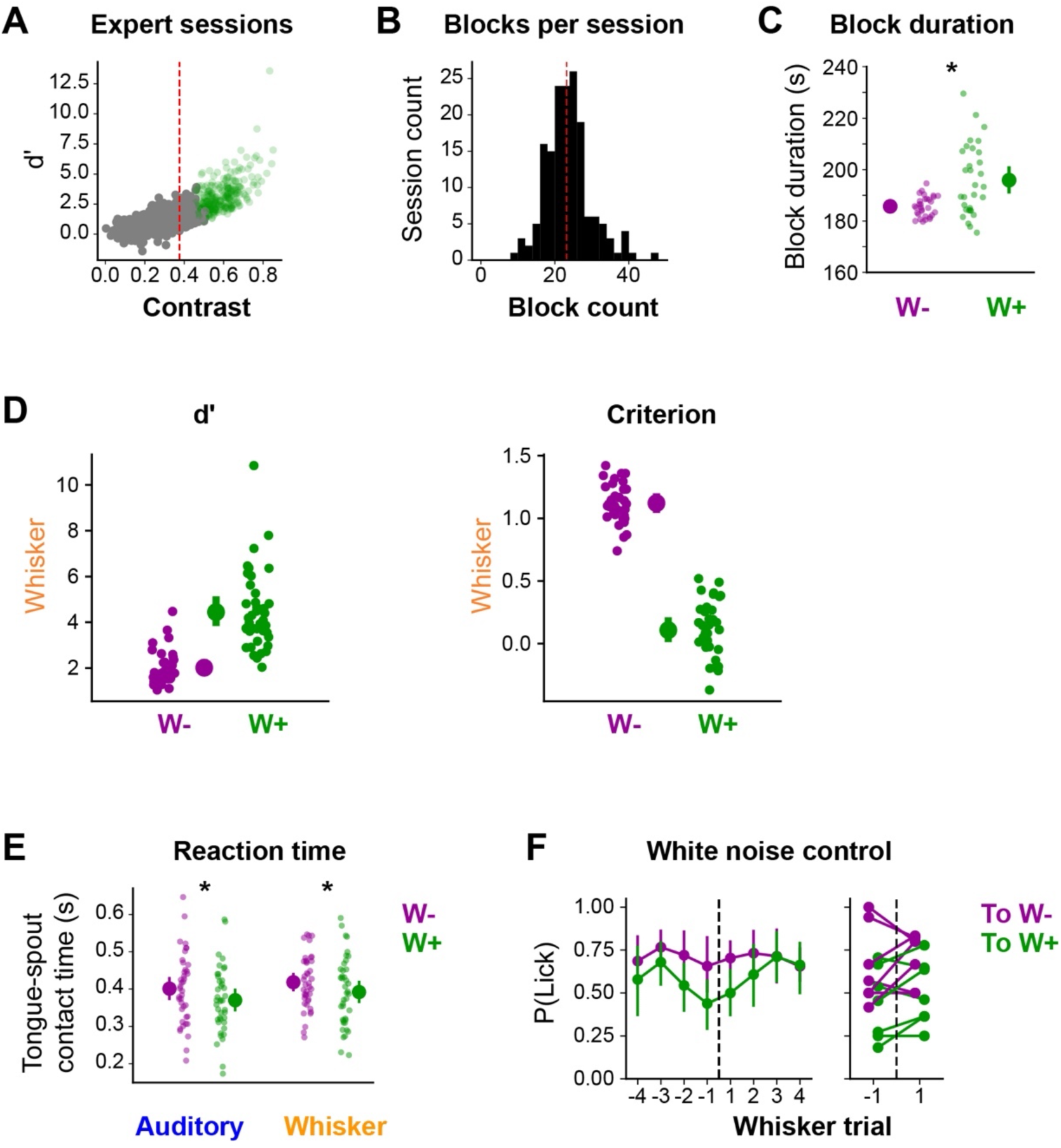
(**A**) Correlation between contrast value used for selection of expert sessions and discriminability index d’ between W- and W+ contexts. Each dot represents a session. The dashed red line represents the threshold for the selection of expert sessions. Sessions with a lower bound 95% confidence interval of mean contrast values (estimated from bootstrapping 10,000 times) above the 0.375 threshold were classified as expert and shown in green. (**B**) Histogram showing the distribution of the total number of blocks in expert sessions. The dashed red line represents the average of the distribution. (**C**) Average duration of single context block. Each dot represents a mouse. Context block durations were averaged within session then further average across sessions to preserved paired data for each mouse. Population average is shown with the 95% confidence interval of the mean. Whisker rewarded context blocks were significantly longer than whisker non-rewarded blocks (W+: 196 ± 13 s, W-: 185 ± 4 s, t-value = 4.3, p-value = 2.0 x 10^-4^). (**D**) Left: Discriminability index d’ between catch and whisker trials in W- and W+ contexts. Right: decision criterion between catch and whisker trials in W- and W+ contexts. Each dot represents a mouse. (**E**) Average spout contact time in response to auditory and whisker trials in W- and W+ contexts showing slightly faster response in the W+ context compared to the W- context for both auditory and whisker trials (Auditory: W+: 370 ± 9 ms, W-: 400 ± 9 ms, t-value = -8.7, p-value = 1.8 x 10^-10^, Whisker: W+: 392 ± 9 ms, W-: 418 ± 9 ms, t-value = -4, p-value = 6.0 x 10^-4^). (**F**) Left: average lick probability in response to whisker trials centred on block transition when contextual backgrounds are replaced by constant white noise. Right: same centre on first trials around transition. Showing absence of context modulated response to the whisker stimulus when context is not explicitly and continuously given to the mice.

**Figure 1 – figure supplement 2.**
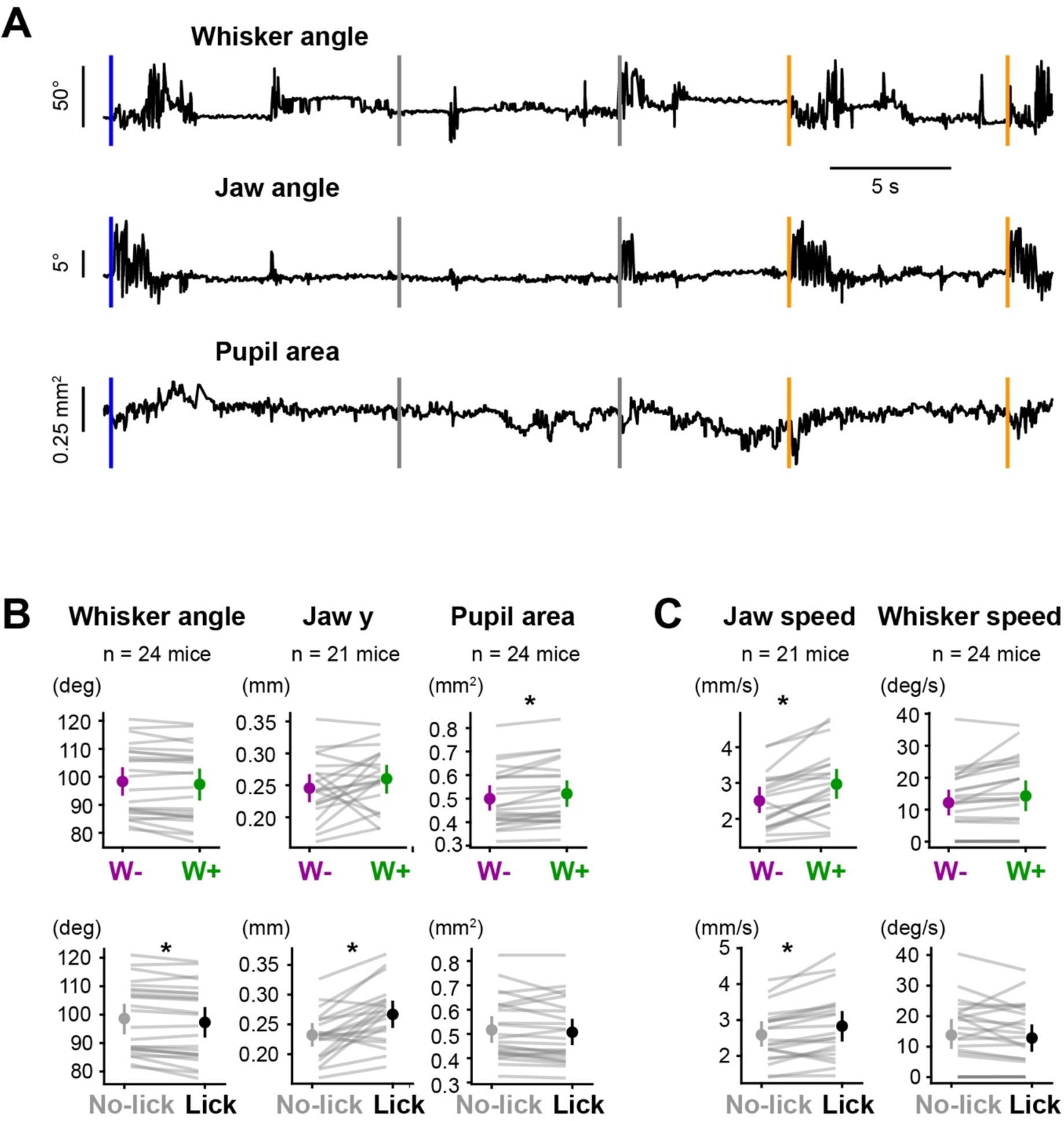
(**A**) Example traces for whisker angle, jaw angle and pupil area. Vertical lines represent trial onsets: auditory, blue; catch, grey; and whisker, orange. (**B**) Tracking of face posture (whisker angle, jaw opening and pupil area) during quiet window preceding whisker trials sorted by context (top row) or by trial outcome (bottom row). Pupil area was significantly modulated by context with larger values in the W+ context compared to the W- context (W+: 0.52 ± 0.13 mm^2^, W-: 0.50 ± 0.12 mm^2^, t-value = 3.7, p-value = 0.023). Whisker angle and jaw opening were modulated by trial outcome with larger values in the baseline preceding lick for jaw opening (Lick: 0.27 ± 0.05 mm, No-Lick-: 0.23 ± 0.04 mm, t-value = 4.6, p-value = 0.0035), and smaller values in the baseline preceding lick for whisker angle (Lick: 97.3 ± 13 °, No-Lick-: 98.6 ± 12 °, t-value = -4, p-value = 0.0103). (**C**) Same as G, but for movement features (jaw speed and whisker speed). Jaw speed was significantly higher in the W+ context compared to the W- context (W+: 3.0 ± 1.0 mm/s, W-: 2.5 ± 0.8 mm/s, t-value = 5.1, p-value = 0.0011) and before Lick compared to No-Lick (Lick: 2.8 ± 0.1 mm/s, No-Lick: 2.6 ± 0.7 mm/s, t-value = -3.7, p-value = 0.0254).

**Figure 1 – figure supplement 3.**
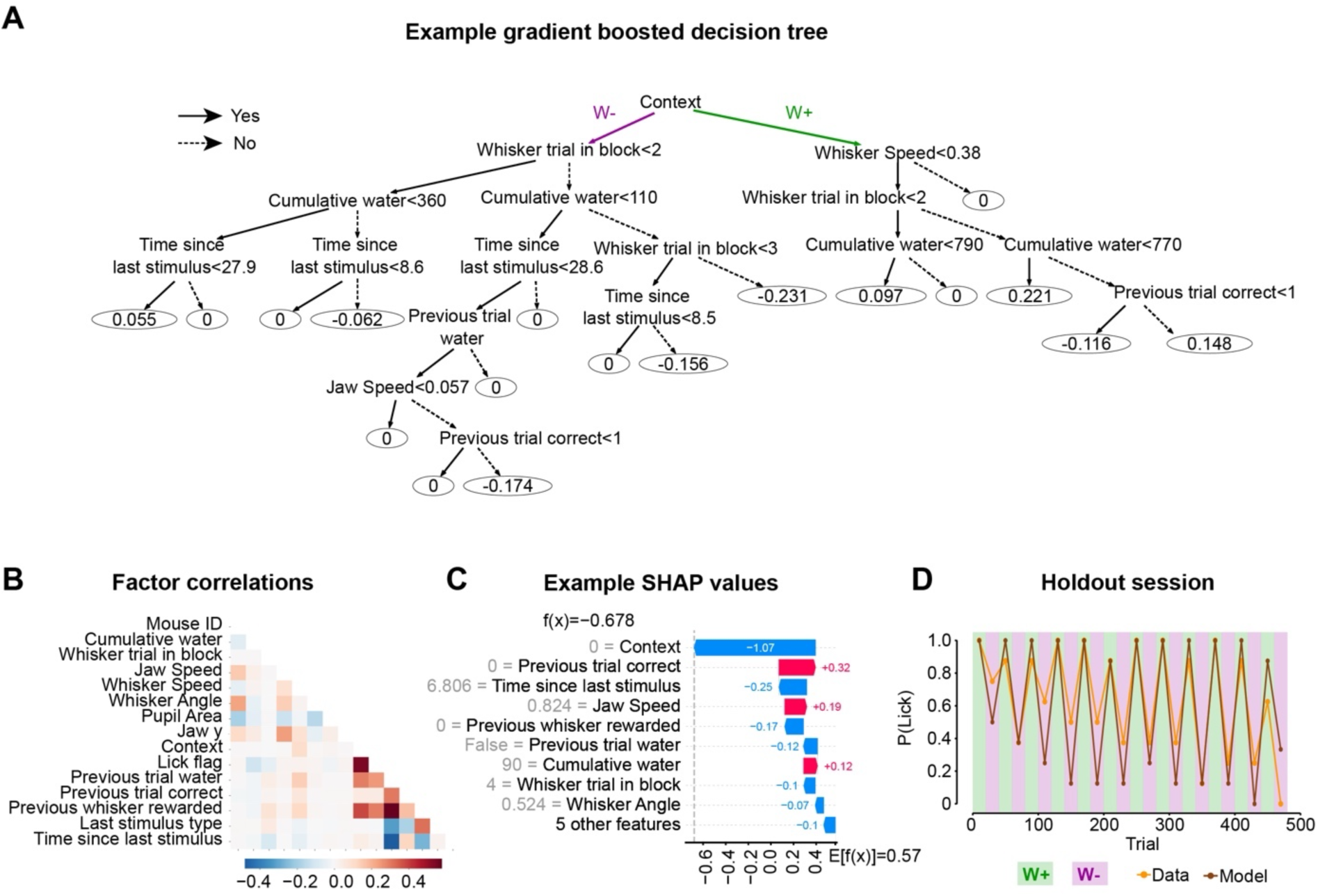
(**A**) Example tree extracted from the gradient boosted tree model showing the impact of parameter values on model decision. Each node of the tree corresponds to a decision step at which a parameter value is used to follow one branch or the other. Positive values in the final leaf tend to deviate the model toward predicting a lick whereas negative values tend to deviate it toward predicting no lick. (**B**) Correlation matrix between the features used to train the behavioral model showing that the model parameters are decorrelated. (**C**) SHAP values for one example trial showing the impact of the parameter value (in grey) on the model prediction for this given trial. Positive values (in red) tend to deviate the model toward predicting a lick whereas negative values (in blue) tend to deviate it toward predicting no lick. (**D**) Performance of the model in one session held out from the training set.

**Figure 2 – figure supplement 1.**
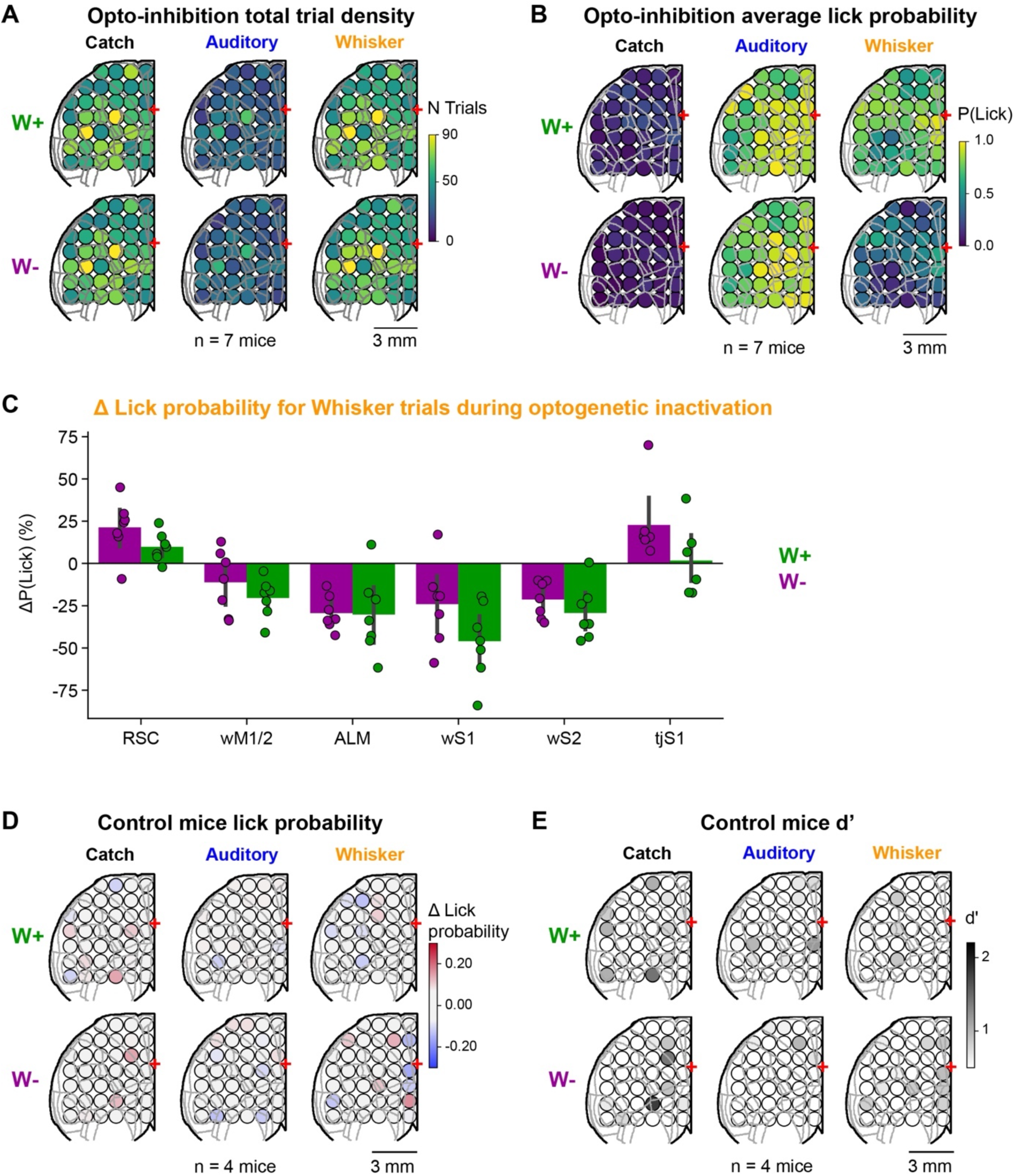
(**A**) Grid showing the total number repetitions of inhibition for each trial type and both contexts at each target location summed over the 7 mice. (**B**) Grid showing the absolute lick probability in catch, auditory and whisker trials in W- and W+ contexts. For each mouse we obtained a grid representation of the lick probability in response to each trial type in both contexts and further averaged these grids over mice. (**C**) Change in lick probability in optogenetic trials compare to control for a selection of cortical regions for the whisker rewarded (W+ green) and whisker non-rewarded (W- magenta) contexts. Each dot represents a mouse, error bars show the 95^th^ confidence interval of the population mean. (**D**) Grid inhibition results in control mice not expressing ChR2. Grid map showing change in lick probability in response to catch, auditory and whisker trials in W- and W+ contexts during optogenetic inactivation of the target area. Changes in lick probability were computed relative to lick probability in the corresponding trial type for each context in control stimulation trials, where the optogenetic laser beam is directed out of the brain. (**E**) Quantification of the optogenetic effect by computing d’ between the distribution of ΔLick probability in stim trials with a null distribution computed by shuffling all trials 1,000 times.

**Figure 2 – figure supplement 2.**
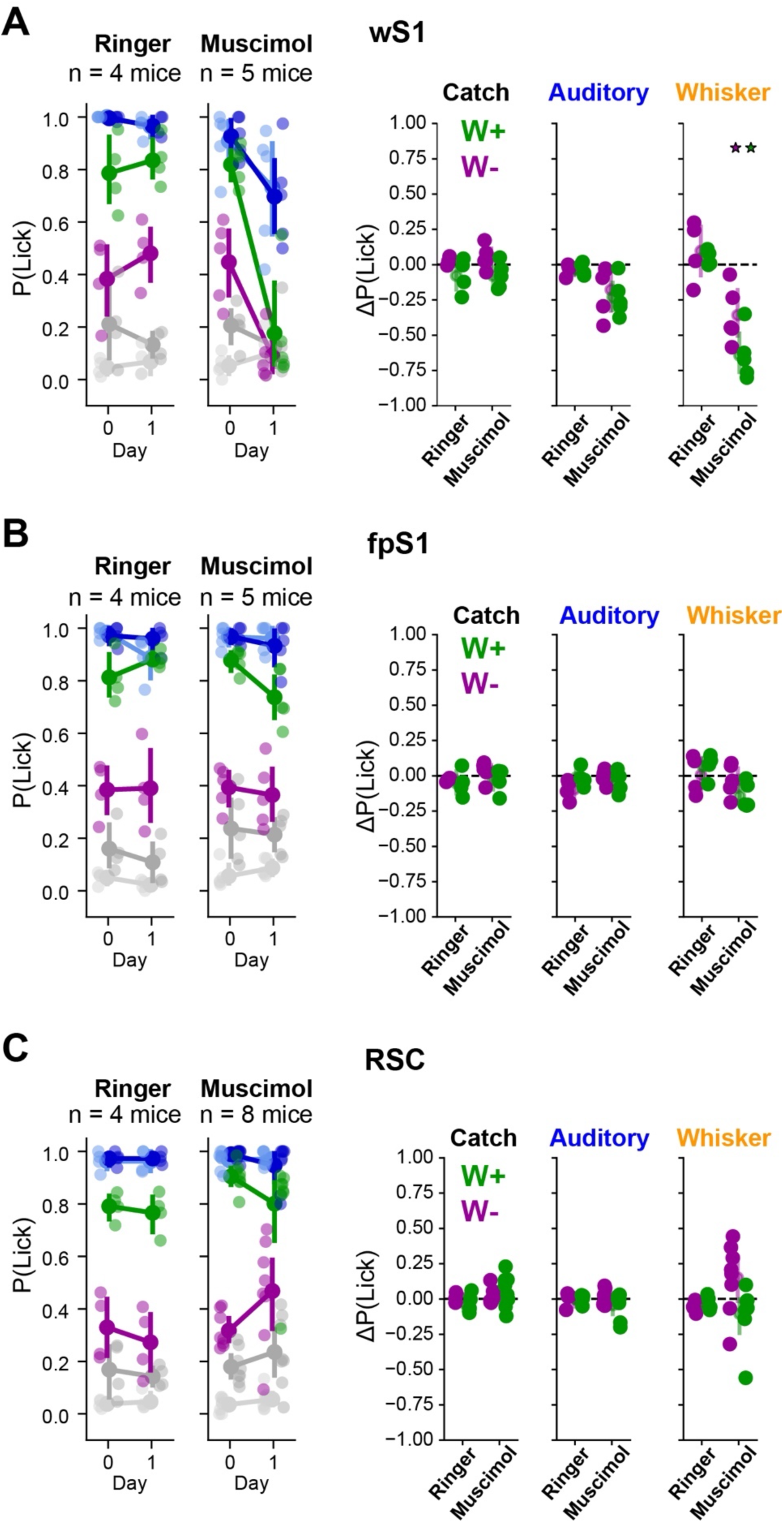
(**A**) Effect of wS1 muscimol or Ringer injection on task execution. Mice were trained until they reliably performed at expert level and on the test day, injections were performed just before the behavior session. Left: lick probability in response to catch, auditory and whisker trials in W- and W+ contexts in baseline day 0 and in day 1 with either Ringer or muscimol injection. Right: change in lick probability in response to catch, auditory and whisker trials in W- and W+ contexts relative to the baseline day. Muscimol injection in wS1 significantly reduced lick probability in response to whisker trials in W- and W+ contexts (hierarchical bootstrapping, p-value < 1.0 x 10^-5^). (**B**) Same as in A, but for injection into the forepaw primary somatosensory cortex (fpS1), showing no significant effect, thus serving as an injection control. (**C**) Same as A, but for RSC inactivation, which leads to a non-significant positive trend in lick probability for whisker trials in the W- context (hierarchical bootstrapping, p-value = 0.059).

**Figure 2 – figure supplement 3.**
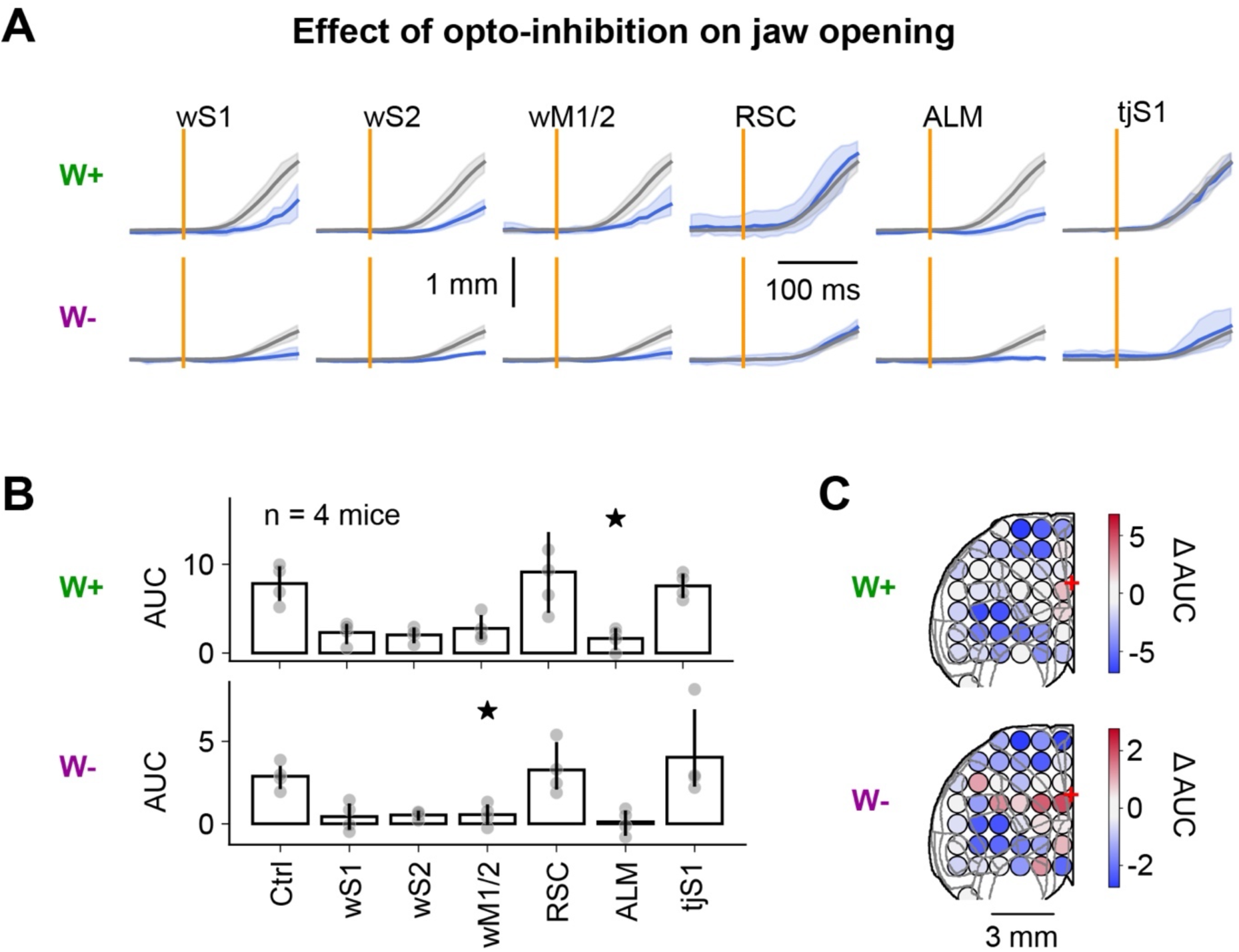
(**A**) Traces of jaw opening aligned on whisker trials for control trials (grey) or optogenetic trials with inactivation of the target area (blue) in W+ and W- blocks. Each dot represents a mouse that was imaged simultaneously with optogenetic inactivation (4 out of 7 mice from the results presented in Figure 2). (**B**) Quantification of the area under the curve in the 150 ms following trial onset for control (first bar) and optogenetic trials (for the 6 target areas) for W+ (top line) and W- (bottom line) contexts. Stars indicate p < 0.05 when comparing the AUC for the optogenetic inactivation of a target area to the ‘Ctrl’ AUC (related t-test corrected for multiple comparisons). **(C)** Grid representation of the average optogenetic effect (change in the area under the curve) on jaw opening compared to control trials.

**Figure 3 – figure supplement 1.**
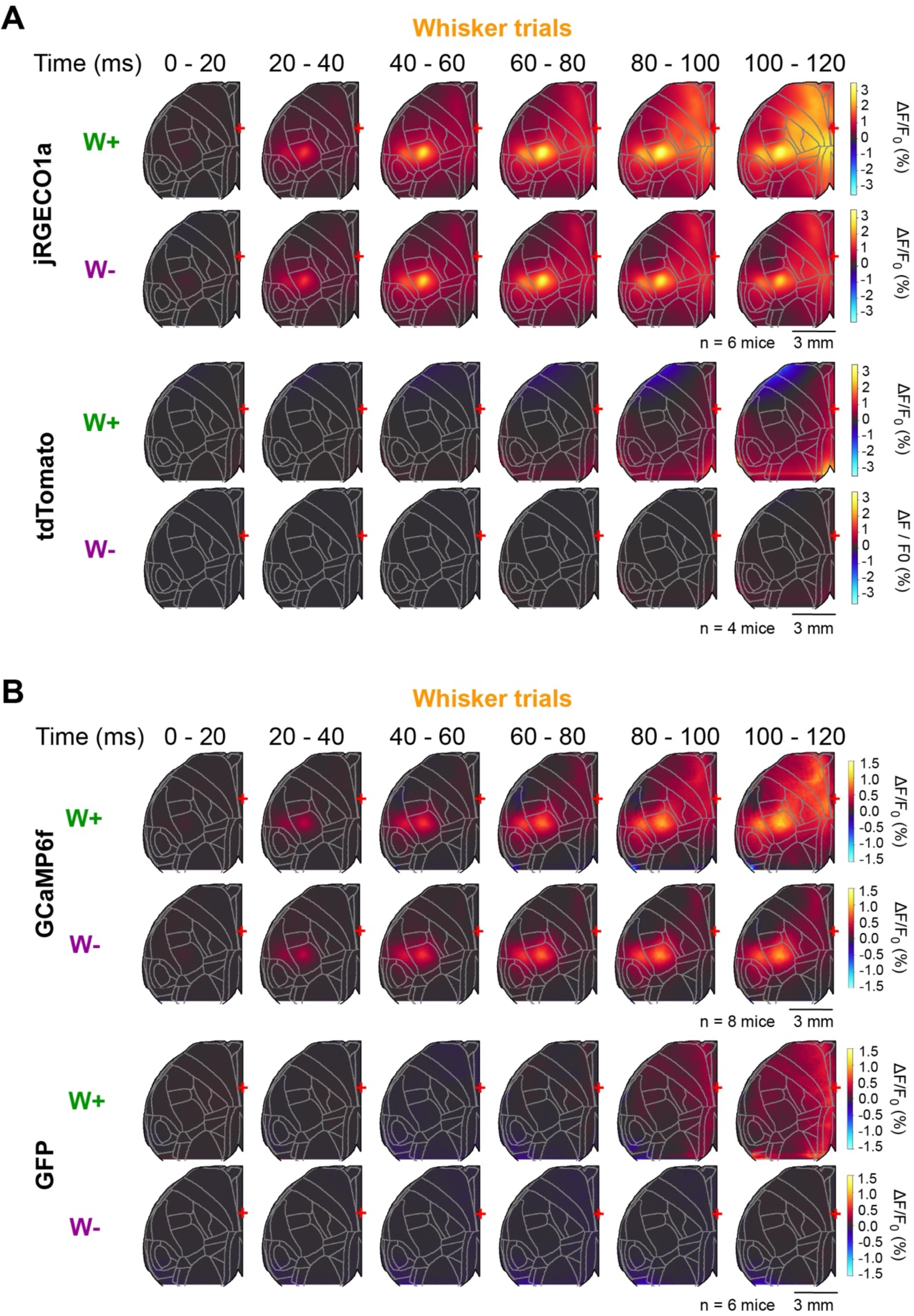
(**A**) Image series showing evolution of cortical dynamics following whisker trials in W+ and W- contexts in jRGECO1a (data duplicated from Figure 3F to facilitate direct comparison) and tdTomato control mice imaged under identical conditions. Early activation of primary and secondary whisker associated somatosensory cortices and propagation of sensory evoked activity were observed only in jRGCO1a expressing mice. (**B**) Same as A, but for GCaMP6f vs GFP mice. Sensory-evoked activity patterns in GCaMP6f-expressing mice were similar to jRGECO1a-expressing mice and not present in GFP-expressing control mice.

**Figure 3 – figure supplement 2.**
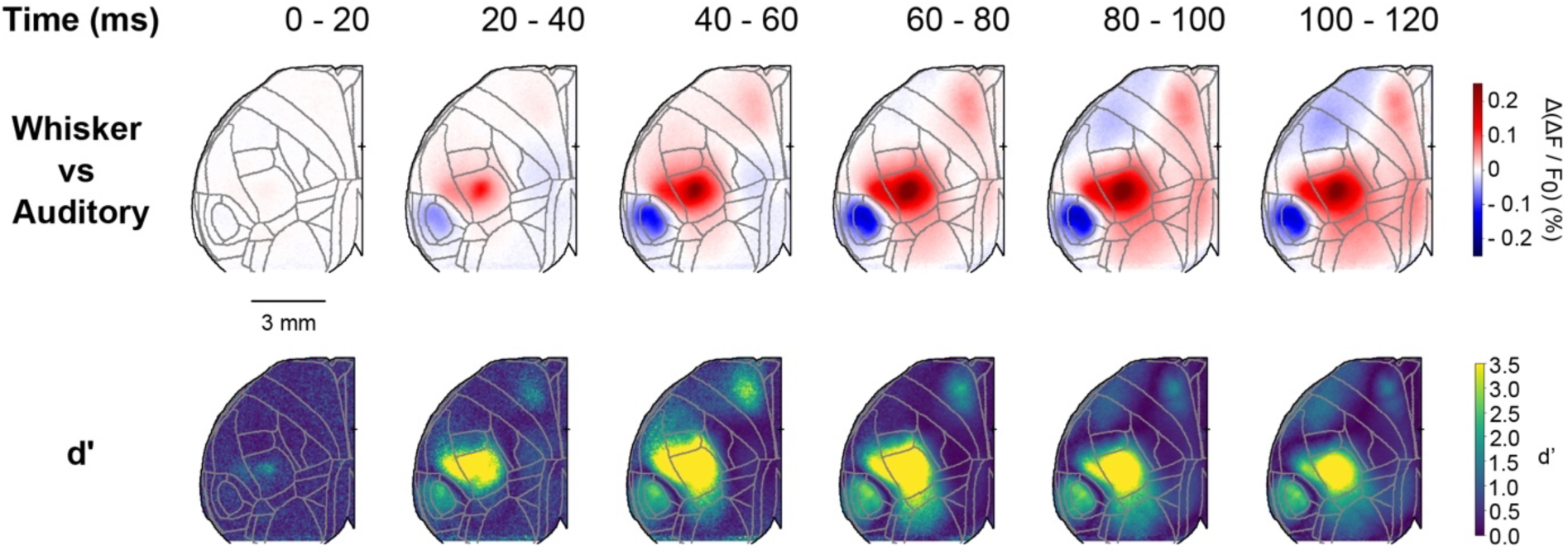
Top: Image series showing evolution of differential cortical dynamics between whisker hit and auditory hit trials in the W+ context (Red, higher activity in whisker hit trials; Blue, higher activity in auditory hit trials). Bottom: discriminability index d’ associated with the mean difference.

**Figure 3 – figure supplement 3.**
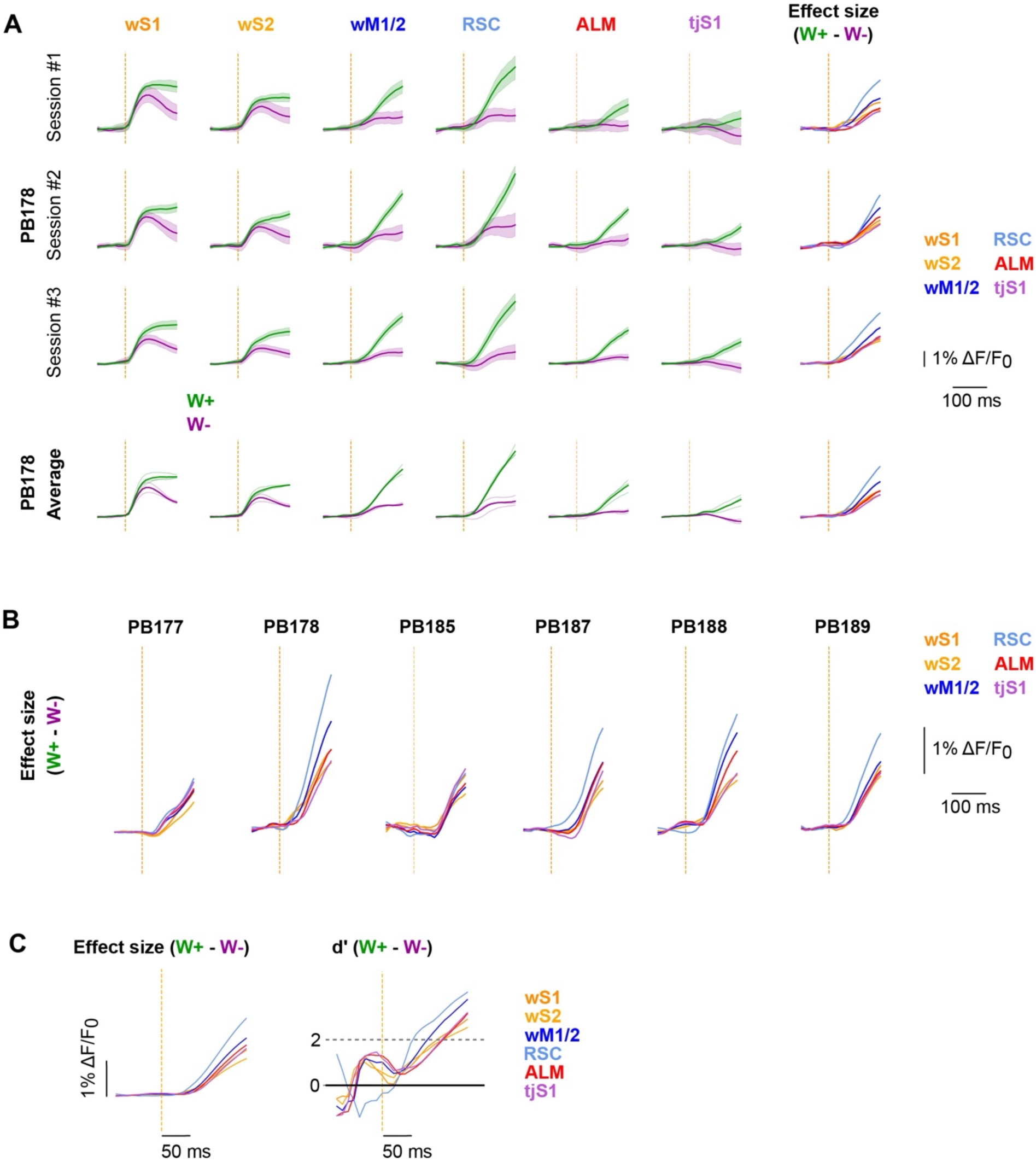
(**A**) Stimulus aligned responses of wS1, wS2, wM1/2, RSC, ALM and tjS1 for correct whisker trials in W- (magenta) and W+ (green) contexts. The first three rows show session results for one mouse (PB178) where shaded area represents the 95% confidence interval around the trial average. The bottom row shows the mouse average resulting from averaging session averaged data in thick lines, each thin line represents a session average. The rightmost column shows the main effect between context defined by the difference of the W- and W+ context means either at the session level (first three plots) or at the mouse average level (fourth plot). Each color represents a selected cortical area. (**B**) Difference of mouse average response of wS1, wS2, wM1/2, RSC, ALM and tjS1 between W- and W+ contexts. (**C**) Left: Difference of population average response of wS1, wS2, wM1/2, RSC, ALM and tjS1 between W- and W+ contexts. Right: Over time discriminability index between context computed from the population (*i.e.*, variance considered between animals). The dashed grey line shows the threshold of 2 used for ranking discriminability between cortical areas in Figure 3H.

**Figure 3 – figure supplement 4.**
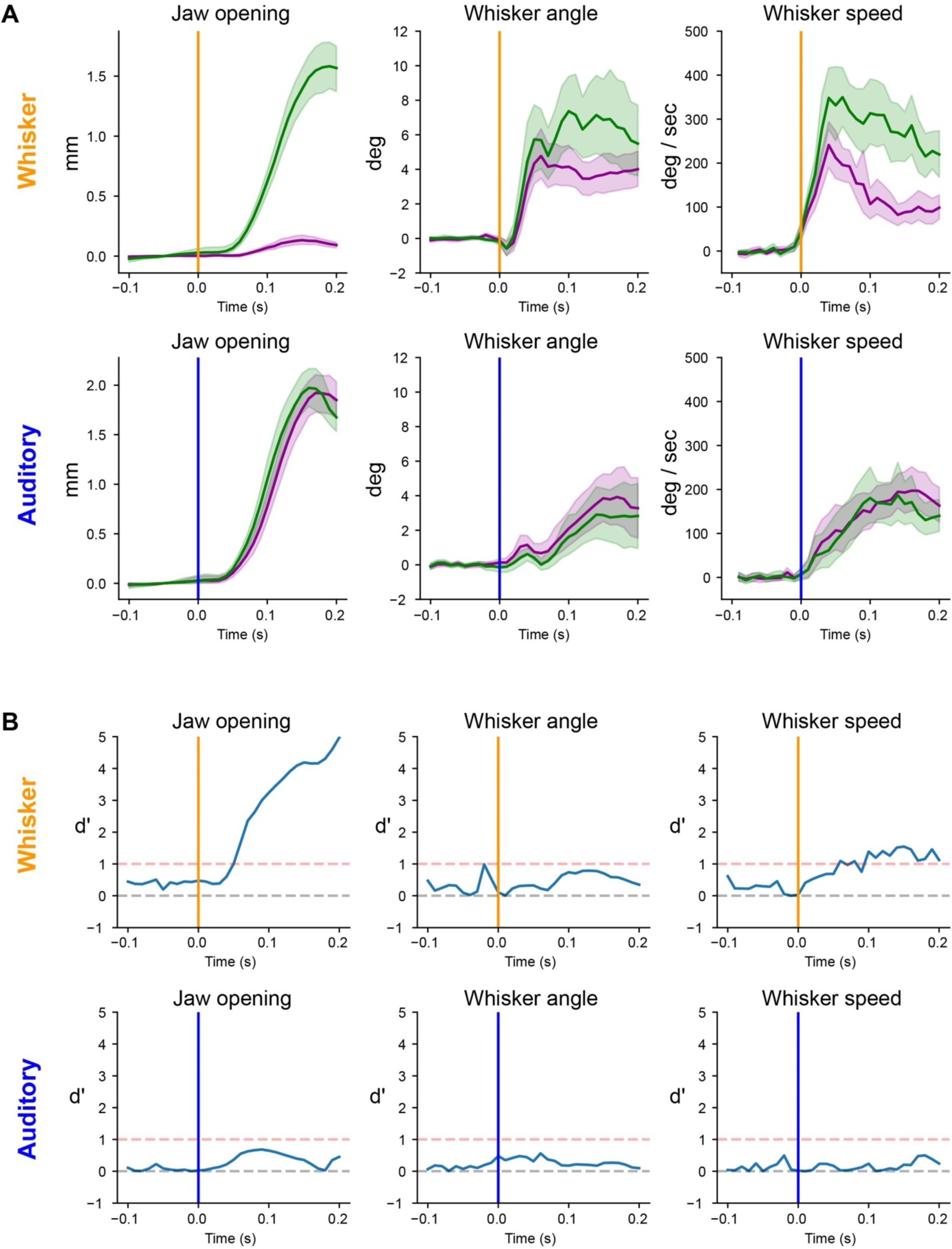
(**A**) Stimulus aligned jaw opening, whisker angle and whisker speed in W+ (green) and W- (magenta) contexts for correct whisker trials (top row) and correct auditory trials (bottom row) (N=24 mice). (**B**) Stimulus aligned discriminability index d’ between W+ and W- contexts for jaw opening, whisker angle and whisker speed for correct whisker trials (top row) and correct auditory trials (bottom row).

**Figure 3 – figure supplement 5.**
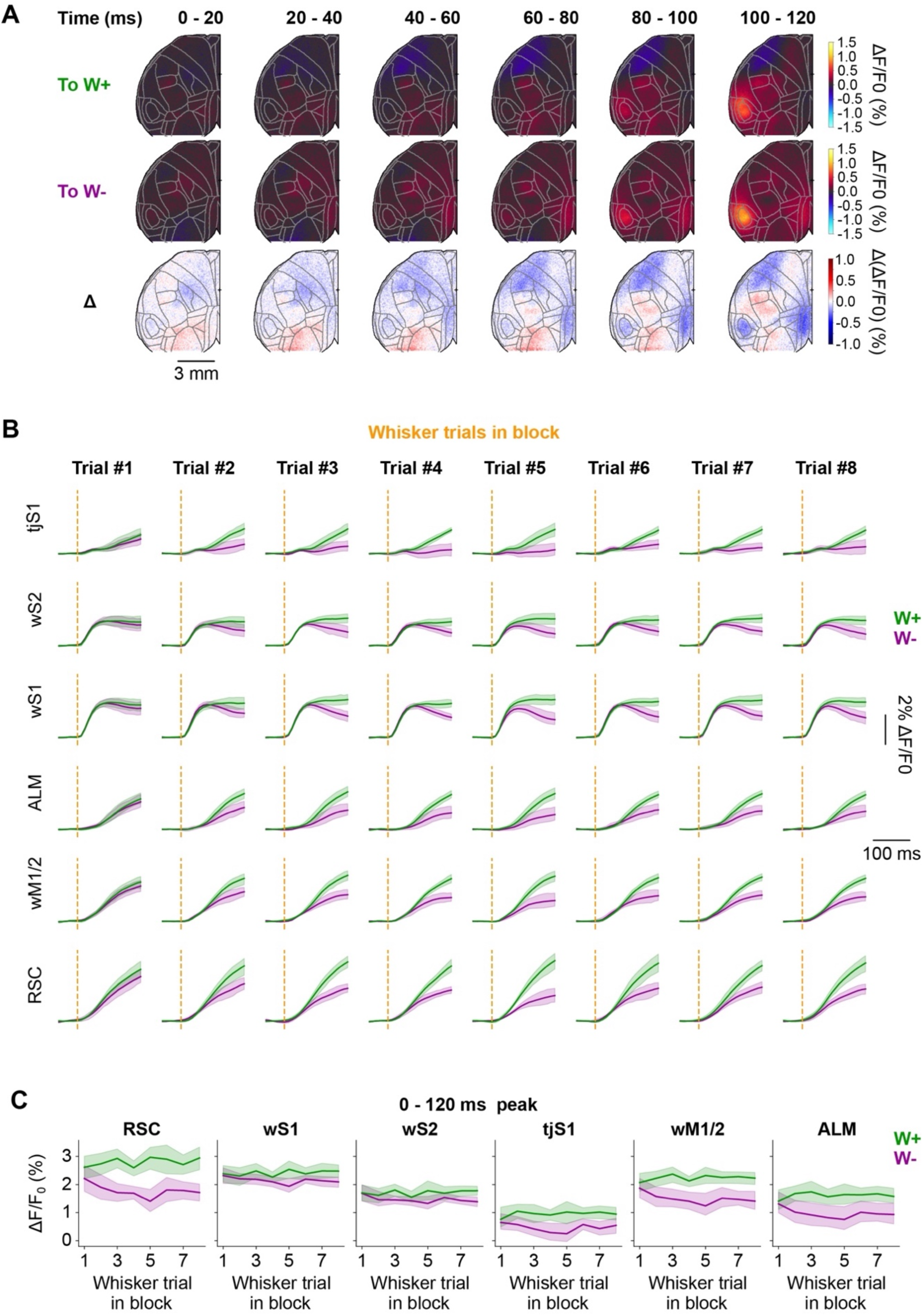
(**A**) Image series showing the evolution of cortical dynamics following the transition to the W+ context (first row), to the W- context (second row), and the difference between activity maps in the W- and W+ contexts (third row). (**B**) Average time-course of stimulus-evoked activity in all whisker trials in W+ (green) and W- (magenta) contexts of selected target areas according to the trial index within the context block. (**C**) Maximal amplitude of the response measured in the 120 ms following the whisker stimulus for each of the selected target area as a function of whisker trial index within context block.

**Figure 3 – figure supplement 6.**
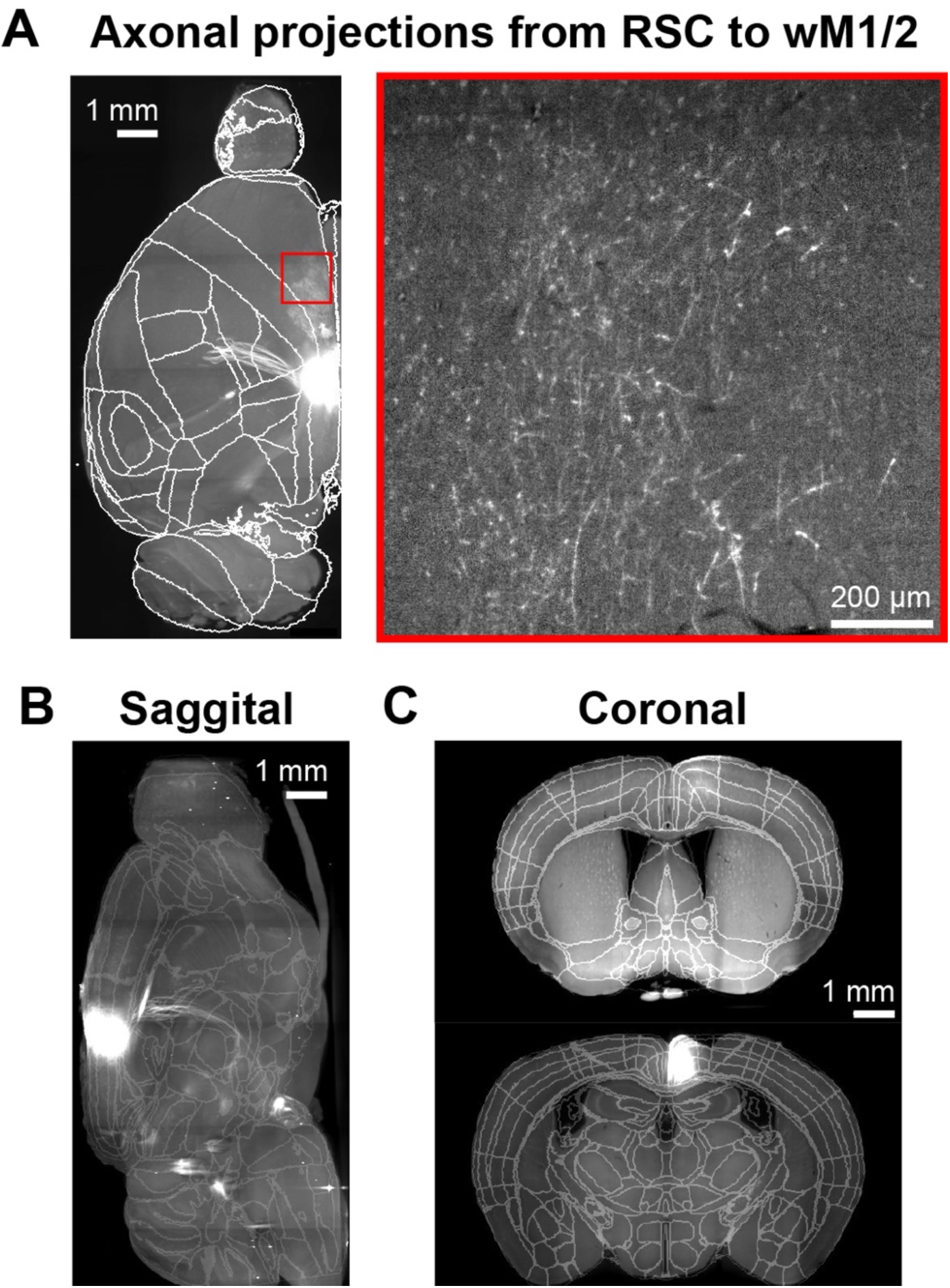
(**A**) To image axonal projections from retrosplenial cortex, mice were injected with an AAV allowing for the expression of membrane bound opsin Chronos fused with GFP (AAV5.Syn.Chronos-GFP.WPRE.bGH). Injections were performed 1.5 mm posterior from bregma and 0.5 mm lateral at 4 depths from the pial surface (-200 µm, -400 µm, -600 µm and -800 µm, 25 nL / site). 6 weeks after injection, animals were perfused and brains extracted to be cleared and immunostained against GFP through iDISCO. We show light sheet imaging sections registered to a mouse brain atlas optimized for light sheet microscopy (Perens et al., 2021) after AAV injection in RSC in an example mouse. Left is a dorsal view of the cleared brain rotated 30 degrees to match the orientation in our widefield imaging, shown as a maximum projection of the most superficial 2 mm of the brain. Right panel shows branching axon innervating wM1/2 at higher resolution, depicted as a single z-plane with thickness of 5.34 μm and x,y voxel size of 1.58 μm. (**B**) Sagittal view around the injection site, shown as a maximum projection of 3 mm from the midline extending laterally. (**C**) Coronal sections of the injection site (bottom) and wM1/2 (top), each shown as a single coronal section resampled to a 20 μm voxel size to improve visualization given the anisotropy of the light sheet images.

**Figure 4 – figure supplement 1.**
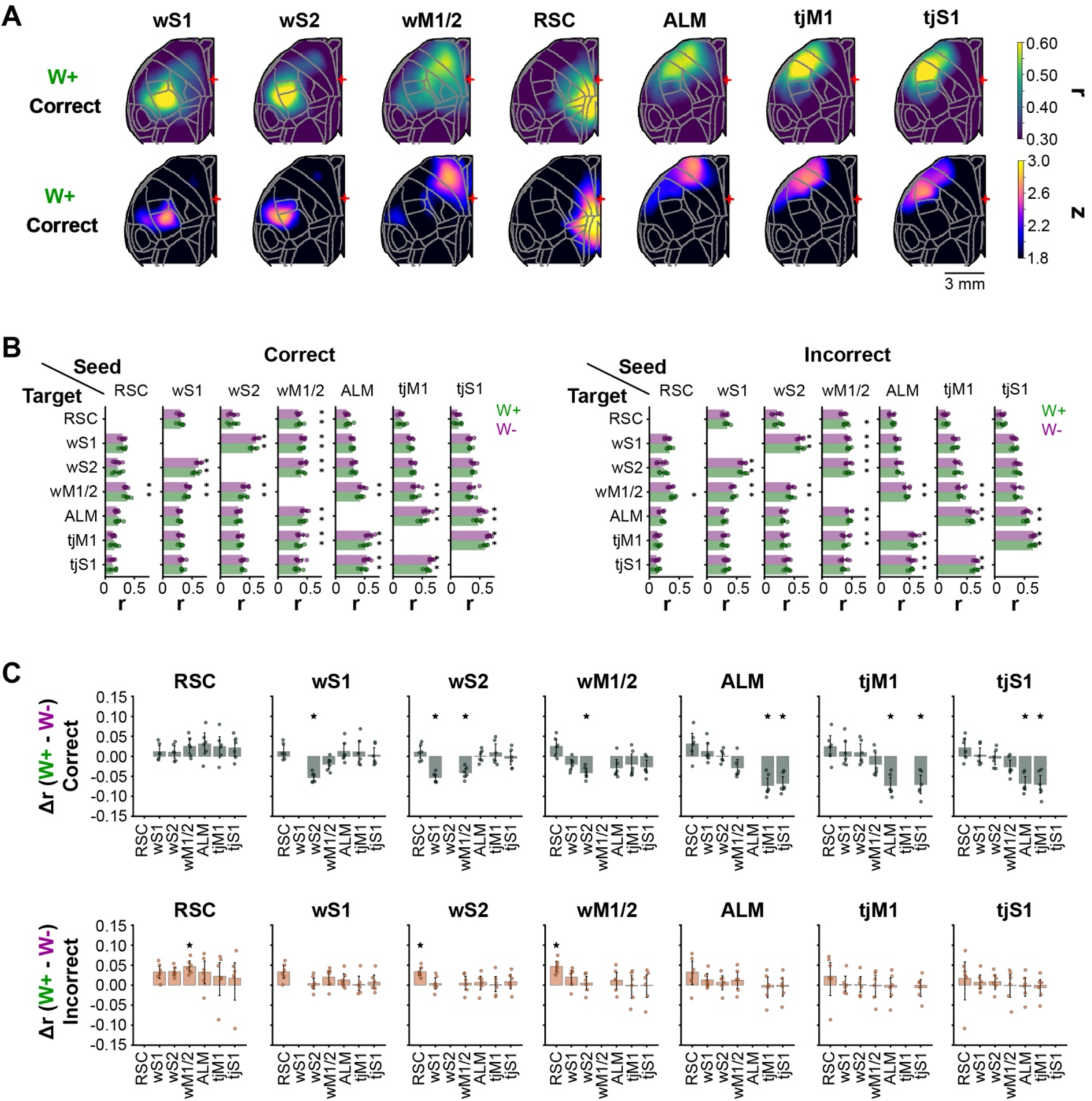
(**A**) Top row: correlation map for wS1, wS2, wM1/2, RSC, ALM, tjM1 and tjS1 seeds in correct whisker trials in the W+ context. Bottom row: After data shuffling, we obtained for each seed and pixel a distribution of correlation value across shuffles. We represented for each seed a map showing the number of standard deviations of the correlation in the observed data compared to shuffled data. (**B**) To investigate correlations between areas, we reduced the correlation maps to grid space. We show as bar plots the correlation between seed and target areas in W- and W+ contexts in correct (left) and incorrect (right) trials. The stars indicate correlation values above 1.8 standard deviations of the shuffled distribution. This allows us to identify significantly ‘connected’ regions, such as wS1 and wS2 or ALM and tjM1 in W+ and W- contexts for both correct and incorrect trials. We used this criterion to obtain the 4 graphs of Figure 4B showing correlation between ‘connected’ regions in W+ and W- contexts for both correct and incorrect trials. (**C**) Correlation changes across contexts between selected seed and target area for correct (top row) and incorrect (bottom row) trials. Stars indicate significantly different correlations in the W- context compared to the W+ context (p <0.05, t-test corrected for multiple comparisons using Bonferroni correction). We used the d’ associated with the amplitude of the context modulation Δ (W+ - W-) or the outcome modulation Δ (Incorrect - Correct) of the correlation for previously identified ‘connected’ regions to build the delta summary graphs shown in Figure 4B.

**Figure 4 – figure supplement 2.**
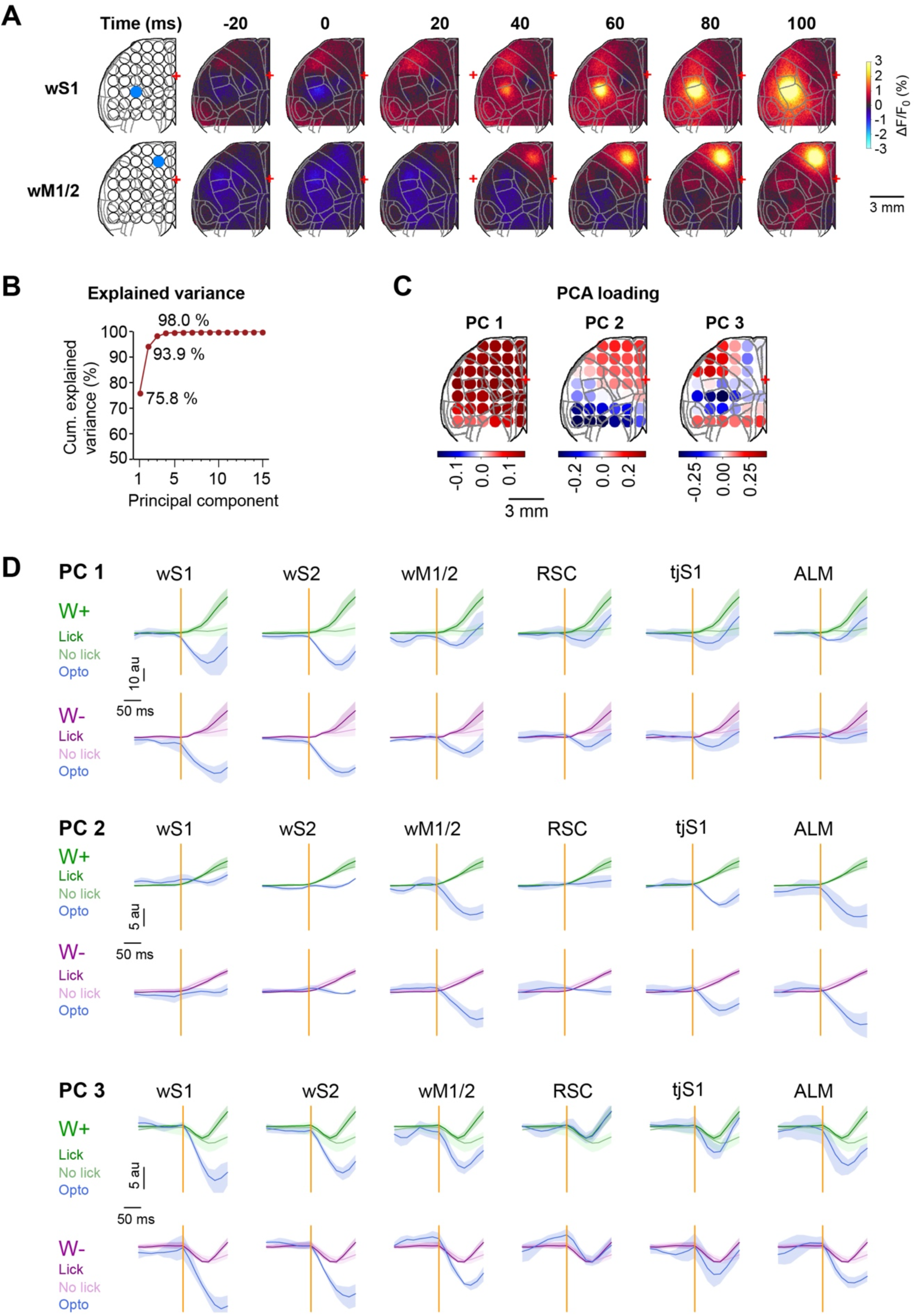
(**A**) Image series showing evolution of cortical dynamics following optogenetic stimulation trials in naïve mice expressing the calcium indicator jRGECO1a - but not the ChR2 opsin - with opto-inhibition laser directed to wS1 (top line) and wM1/2 (bottom line). This control experiment shows a locally restricted jRGECO1a photo-activation effect (jRGECO1a is based on mApple, which shows photoconversion upon blue light stimulation), but no widespread inhibition as observed in mice expressing the ChR2 opsin in inhibitory neurons (Figure 4C). (**B**) Cumulative variance explained by the first 15 components resulting from PCA. (**C**) Grid representation of the loading on the first 3 PCs of each of the grid locations. (**D**) Time course of PC1, PC2 and PC3 in W+ and W- contexts for control and selected optogenetic inhibition points for control trials with lick or no lick and optogenetic trials.

